# The evolutionary processes of bacterial aromatic polyketide ketosynthases

**DOI:** 10.64898/2026.07.16.738910

**Authors:** Xiaoyu Wang, Qiandi Gao, Liangjun Ge, Zhiwei Qin

**Affiliations:** Center for Biological Science and Technology, Advanced Institute of Natural Sciences, Beijing Normal University, Zhuhai, Guangdong, 519087, China

**Keywords:** Natural product, polyketide synthase, evolutionary dynamics, large language model

## Abstract

**Background:** The biosynthesis of bacterial aromatic polyketide polyketides (type II polyketides, T2PKs) employs a single set of catalysts (ketosynthases, KSs or KS_α_, with chain length factors, CLFs or KS_β_) and iteratively assembles a carbon backbone with precise chain length control. Considering the increasing number of T2PKs discovered in laboratory settings, it is necessary to understand the evolution trajectories of KSs and CLFs.

**Results:** We employed our recently developed algorithm, MAAPE, based on large protein language model (PLM) to glean insights into the evolution process of KSs and CLFs. Our findings indicated the evolutionary history of KS and CLF domains from bacterial T2PKSs and identified a shared ancestral cluster (Cluster A), supporting a common origin. Despite structural homology, KSs and CLFs followed distinct evolutionary paths, shaped by coevolution and early horizontal gene transfer.

**Conclusions:** Understanding the evolutionary lineage of these enzymes will illuminate the natural optimization processes of their functions and present opportunities for the rational design of novel polyketides with enhanced efficacy.

## Background

The biosynthesis of bacterial type II polyketide natural products (T2PKs) are catalysed by type II polyketide synthases (T2PKSs). T2PKSs are important because of their significant roles in natural product biosynthesis processes, and they have led to the discovery of numerous medically and industrially relevant compounds [1, 2]. A T2PKS typically consists of a heterodimer of a ketosynthase (KS, or KSα) and a chain length factor (CLF, or KS_β_), an acyl carrier protein (ACP), and a malonyl-CoA:ACP transacylase (MAT). The collective action of the KS, CLF, and ACP, which is referred to as a minimal PKS, initiates polyketide biosynthesis by decarboxylating malonyl-ACP [3]. A KS catalyses the polyketide chain elongation process through the Claisen condensation reactions of acyl-thioester units such as malonyl-CoA and methylmalonyl-CoA, whereas the CLF governs the chain length via C-C bond formation during each iterative cycle [4]. These enzymes generally collectively determine whether to continue the elongation process or terminate the chain after the assembly of a specific number of building blocks. Such far, approximately 166 T2PK families have been identified based on building block counts of 8, 9, 10, 12 and 13, respectively [5].

The current understanding suggests that KSs and CLFs originated from a common ancestor. Following a gene duplication event, CLFs diverged from KSs, driven by evolutionary pressure to fine-tune the polyketide chain length control scheme [6]. While KSs maintained a more conserved evolutionary path, CLFs underwent rapid adaptation, likely in response to environmental pressures that demanded greater polyketide product diversity [7, 8]. Although certain aspects of this evolutionary theory have been supported by experimental data and traditional phylogenetic analyses, they remain largely speculative and require more rigorous validations. A comprehensive verification likely requires the development of novel methodologies and further experimentation.

We recently proposed DeepT2, a classification model for CLFs that builds on sequence embeddings from the protein language model (PLM) ESM-2 [9], which was pretrained on large-scale protein sequence data. Using these embeddings, DeepT2 achieved an F1 score of 0.98 while being trained only on a dataset comprising CLFs with known T2PK scaffold structures and 2566 CLFs with unknown scaffolds. In this ongoing research, we aim to leverage state-of-the-art PLMs to reveal the evolutionary relationships between KSs and CLFs [10]. The intriguing interplay between KSs and CLFs is pivotal because they maintain a delicate balance between chain elongation and termination. These interactions have evolved to ensure precise control over polyketide lengths and compositions, enabling microorganisms to adapt their secondary metabolites for use with various ecological and survival strategies [11].

Here, we report a novel perspective concerning the evolution dynamics between bacterial KSs and CLFs via PLMs. The primary objective of this project is to investigate whether the embeddings (the features of a protein) derived from PLMs can be analyzed and reconstructed to quantify the evolutionary relationships between protein sequences [12]. We employed Modular Assembly Analysis of Protein Embeddings (MAAPE), a PLM-based strategy recently developed by our group, to investigate whether protein features derived from PLM embeddings can elucidate evolutionary relationships [13]. Provided that the evolution process is predictable, biosynthetic engineers can leverage the findings presented herein to rationally design enzymes with tailored functions, thereby creating a versatile array of molecular tools for a wide range of applications.

## Results

### Discrepancy between general phylogenetics and PLM embeddings

Previous phylogenetic analyses conducted at either the gene or gene cluster scale have elucidated the chemical diversity of T2PKs [14, 15]. In this study, our focus lies in employing PLMs to investigate the potential evolutionary history of KSs and CLFs. To ensure a comprehensive understanding, it is imperative to consolidate the earlier findings using the most recently updated database. It is worth noting that that the dataset of 166 KS-CLF pairs was curated from biosynthetic gene clusters with experimentally characterized products, retaining only full-length, non-redundant sequences that could be unambiguously assigned to core aromatic polyketide scaffolds, while excluding clusters with incomplete annotation, ambiguous domain organization, or high sequence redundancy (>90% identity). This compilation will serve as a crucial reference point for our subsequent investigations. Consequently, we meticulously organized a dataset comprising 166 pairs of bacterial KSs and CLFs, whose corresponding chemical structures are known, to perform a large-scale phylogenetic analysis (Supplementary Table S1). As depicted in Figure 1a, the results revealed 2 distinctly divergent clusters of KSs and CLFs, which were generally consistent with those obtained in previous analyses [7]. However, we have not observed any direct evidence regarding the evolution from KSs to CLFs, or vice versa.

Inspired by recent developments in theoretical bioinformatics and aiming to resolve these ambiguities, we developed a novel evolutionary inference approach for the same dataset. Instead of directly analysing information coded by a combination of 20 amino acids, we strived to extract what is called the “Grammar of Life” from protein language-embedded sequences. In this work, we utilized ESM-2, a large-scale PLM that was pretrained on hundreds of millions of protein sequences with up to 15 billion parameters, to capture evolutionary information and structural features at the atomic level [10]; the speed of this model is comparable to those of AlphaFold2 and RoseTTAFold, facilitating a rapid exploration of the structural space of metagenomic proteins [16, 17].

**FIGURE 1.**
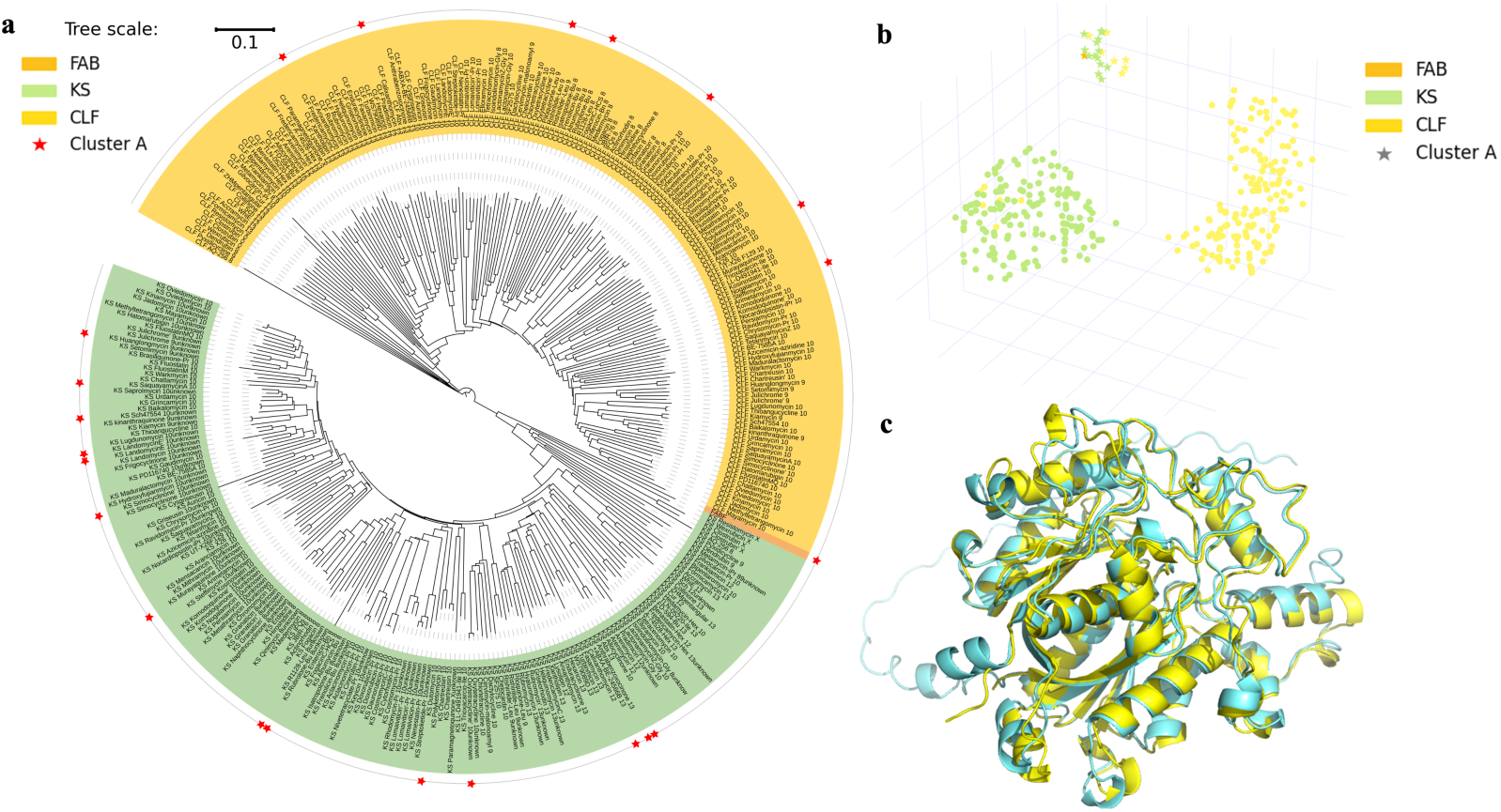
Comparison between a phylogenetic tree constructed from 166 pairs of structurally analogous bacterial KS-CLF proteins and a *Staphylococcus aureus* FABF and a visualization of the UMAP-based dimensionally-reduced ESM-2 embeddings to a 3D version of the same dataset; categories are grouped by colour, and Cluster A nodes are marked by stars. Multiple sequences were aligned via CLUSTALW. (a) The phylogenetic tree was constructed via IQ-TREE, with the LG+F+R8 model selected according to the Bayesian information criterion. (b) Visualization of KS-CLF-FABF embeddings. (c) AlphaFold3 predicted the 3D structures of the KS and CLF domains from actinorhodin T2PKSs; the KSs & CLFs exhibited structural analogy, with an RMSD of 0.993 during structural alignment. This figure was created using BioRender.

Figure 1b shows the results of dimensionally reducing the ESM-2 embeddings of KS and CLF sequences, which align with the phylogenetic analysis, where KS and CLF sequences form distinct clusters due to their substantial sequence divergence (identities below 30%). Notably, we observed several CLF nodes within the KS cluster, and more intriguingly, a distinct cluster of 22 nodes (marked with stars) containing both KS and CLF sequences. This mixed cluster deviates from the pattern observed in the phylogenetic tree, with the *Staphylococcus aureus* FabF sequence, serving as an outgroup, also present within it. Given that bacterial FabF is functionally analogous to KS but highly divergent, we propose this mixed cluster represents ancestral KS and CLF sequences [6, 18]. A detailed list of the members of this ancestral group is presented in Supplementary Table S2 and is referred to as “Cluster A” in this work.

Upon mapping the proteins obtained from Cluster A onto the phylogenetic tree in Figure 1a, we observed a dispersed distribution, indicating that the PLM captured extra information from the sequence analysis alone. A comparative analysis of the tertiary structures of the KS and CLF proteins derived from the same BGC revealed remarkably high structural similarity, with RMSD values less than 1 for structural alignments. Figure 1c shows the structural alignment of the KS and CLF determined from the actinorhodin biosynthetic pathway, which revealed high structural similarity despite their low sequence identity, which is a hallmark of structural homology. This observation suggests that while these proteins have undergone substantial sequence divergence, they have maintained their structures and functions throughout their respective evolutionary trajectories.

These findings demonstrate the limitations of evolutionary analyses based solely on sequence alignments for structurally homologous proteins such as KSs and CLFs and highlight the importance of employing PLMs for evolutionary analysis purposes, particularly in cases where traditional sequence-based methods may be insufficient for elucidating the complex evolutionary histories of structurally conserved but sequentially divergent proteins. Consequently, we applied MAPPE to explore the evolutionary trajectories of this functionally intriguing polyketide synthase family, providing fresh insights into their evolutionary dynamics and functional refinement.

### Evolution trajectories of KSs and CLFs

#### 1. Cluster A as the ancestral group

We first used MAAPE to obtain the embeddings of 166 known T2PKS KS/CLF pairs. While our results were largely consistent with those of classic methods, they offer more granular insights into evolutionary trajectories and provide refined ancestral state predictions. By integrating edge weights and directional information from the embedding co-occurrence matrix into the KNN network, we confirmed the predicted root position of Cluster A (Figure 2a). The root node identified through the diffusion analysis (located within this region), combined with in-degree and out-degree analyses of the directed KNN graph, collectively support this validation[12] (Figure 2a; the displayed numbers correspond to the sequence IDs within Cluster A).

**FIGURE 2.**
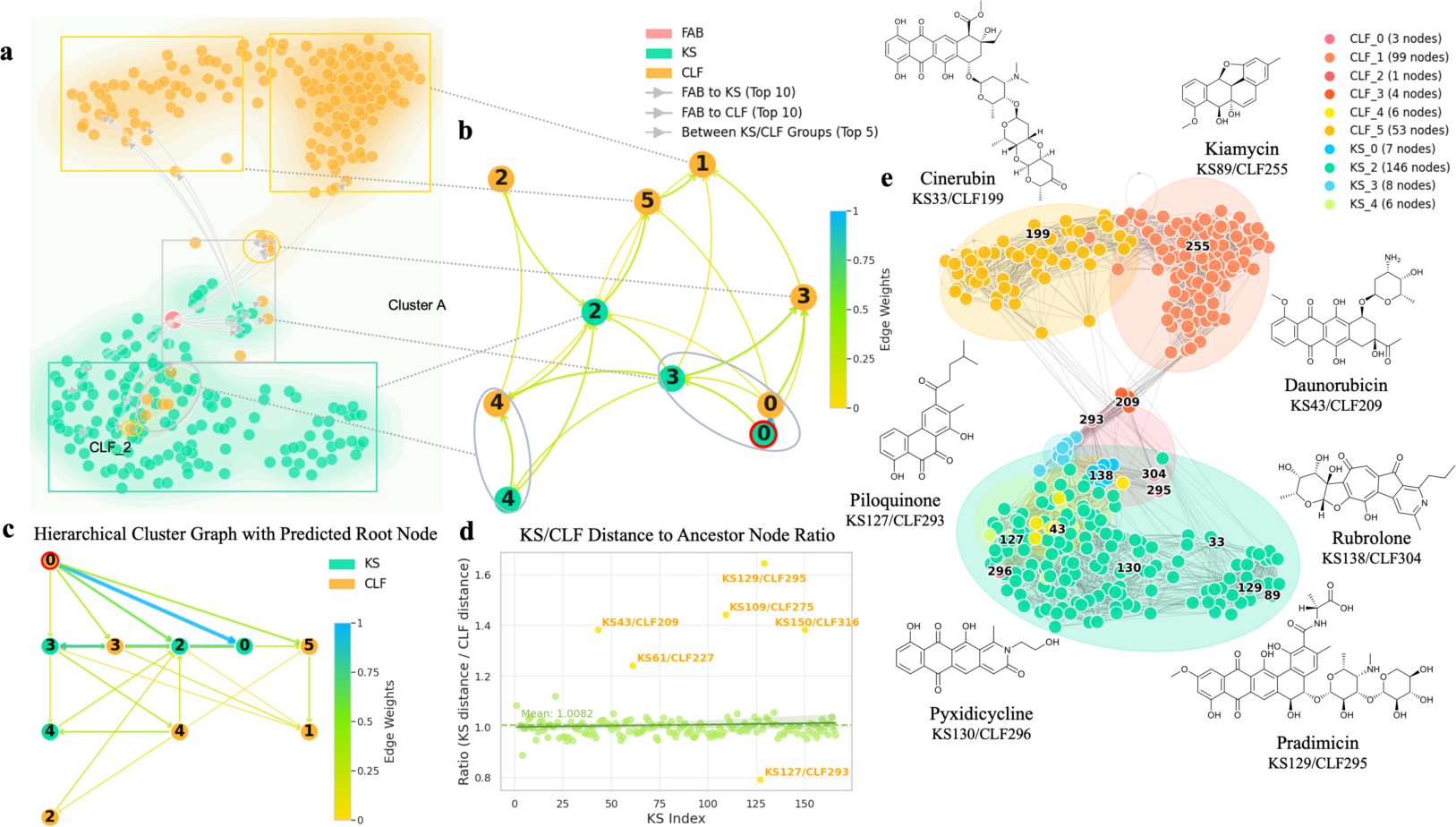
Evolutionary trajectories of known KS/CLF protein sequences determined by the MAAPE algorithm with selected product structure annotations. (a) This network was arranged based on the Euclidean distances between the vector representations of the 166 KS/CLF sequences obtained through a KNN similarity graph built upon ESM-2 embeddings and overlaid with edge directions derived from the MAAPE method, only the top 10 highest-weighted edges from FAB to KS/CLF clusters and the top 5 between KS and CLF clusters are displayed. Nodes with smaller Euclidean distances tended to cluster together in the KNN similarity network. Edges connecting the nodes were directionality assigned based on the calculated assembly directions of the PKS modules, revealing how horizontal transfer and assembly events gave rise to new enzymes during the T2PKS evolution process. Arrows indicating stronger signals for module transfer and assembly directions. The node of Cluster A is shown in the middle area. (b) The nodes were aggregated according to their KS or CLF groupings and the edge-bundled version of Figure 2a, and the thicknesses of the edges are reflected by the resulting edge weights. The KS sequences were clustered into 4 groups, the CLF sequences were clustered into 6 subgroups, and the subgroups were distinguished according to their colours and numbers. CLF_0 was predicted to be the root group via the diffusion analysis, so it is circled in red in both Figure 2a and Figure 2b. KS_0 and KS_3, as highlighted with green circles in Figure 2a, showed close proximity to CLF_0, suggesting close relationships with the root of the network. A cluster of CLF nodes was found in the KS regions, and these nodes were grouped as CLF_2&4 and highlighted with orange circles in Figure 2a. (c) Rearrangement of Figure 2b on the basis of hierarchical relationships. (d) KS/CLF distances to the anchor node. The ratios of the distances from the KSs and CLFs to their ancestor nodes were plotted against the KS index sequence. Most data points clustered around a ratio of 1, indicating coevolution between KSs and CLFs. Yellow points highlight outliers that significantly deviated from the average, and they are labelled with their KS and CLF indices. (e) Representative product structures for selected KS/CLF pairs from different clusters, with their associated product structures and KS/CLF index displayed around the network periphery. Various groups are highlighted with distinct colored backgrounds corresponding to their cluster assignments. This visualization demonstrates the functional diversity of secondary metabolites produced by different evolutionary clusters. Specific cluster assignments include: Cinerubin (KS_2/CLF_5, both from majority clusters), Pradimicin and Piloguinone (both KS_2/CLF_0, representing KS majority group/CLF Cluster A and identified as outliers in KS/CLF distance ratio analysis) and Pyxidicycline (KS_4/CLF_2, non-oxidative T2PKS). This figure was created using BioRender.

Cluster A comprises 22 sequences, including FabF, 7 CLF nodes, and 14 KS nodes - double the number of CLF nodes. Given that KSs are presumed to be more primitive than CLFs are, the dominance of KSs in this cluster aligns with this hypothesis. However, the topological structure within this cluster presented a reversal signal, with 3 CLF nodes predicted to be roots. This finding alone cannot overturn the conventional view, as biases may have existed in our product-oriented dataset of 166 sequences. Additionally, the observed strong ancestral signal in the MAAPE analysis could be influenced by the gene losses of early CLF sequences or neofunctionalization events.

Notably, except for one KS-CLF pair associated with Rubrolone-Bu, nearly all nodes in Cluster A are unpaired. This suggests that KSs and CLFs may have initially originated from a single gene or protein complex with a common ancestry, which subsequently diverged into distinct entities. The separation of KSs and CLFs into independent components might have occurred during a single evolutionary event, after which they evolved independently. This could explain the lack of pairing within this ancestral cluster.

#### 2. Evolution of the KS and CLF domains

In Figure 2a, we observed a Z-shaped pattern, with Cluster A dispersed along the central arm, whereas the other KS and CLF nodes diverged into two distinct symmetric clusters. To better understand their evolutionary trajectories, we grouped the nodes on the basis of their KS and CLF categories and applied edge bundling to produce a condensed visualization [19] (Figure 2b). This updated figure illustrates how the KS and CLF sequences originated from a common ancestor (CLF_0 & KS_0) and subsequently evolved independently. The node lists for these aggregated clusters are provided in Supplementary Table S3, and a hierarchically sorted version of this figure is presented in Figure 2c. Furthermore, the inter-cluster edge weights and directions after clustering analysis are recorded in Supplementary Table S4.

CLF_0, which was predicted to be the root group, is highlighted with a red stroke and corresponds to the nodes circled in red in Figures 2b and 2c. The locations of the KS_0, KS_2 & KS_3 groups, which are near CLF_0, suggest not only their structural and functional similarity but also their close evolutionary relationship to the ancestral state. Excluding the ancestral node of Cluster A, the KS and CLF sequences clearly diverged into two distinct groups, indicating that their descendant lineages differentiated from a common ancestor into the KS_2 group and the CLF_1&5 group, respectively. Notably, KS_2 and CLF_5 are directly connected to the root by edges with large weights, indicating strong gene transfer events. While CLF_1 is positioned on the same side of the Z shape as CLF_5 in Figure 2a, the hierarchical diagram in Figure 2c reveals that its evolutionary relationship is posterior to CLF_5, receiving gene transfers via two distinct paths: CLF_0-KS_3-CLF_1 and CLF_0-CLF_3-CLF_1. This analysis provides a broad overview of how the KS and CLF genes evolved from Cluster A into two independently diverging clusters. Nodes that are not explicitly mentioned within these key clusters may have contributed specific sequence modules to the primary group or represented evolutionary attempts to adapt to environmental pressures.

#### 3. CLF evolution trend for nonoxidative T2PKs

During the MAPPE analysis, we observed that non-oxidative T2PKs form a relatively isolated group in Figure 2. The primary mechanism for aromatic ring formation in non-oxidative T2PKs bypasses oxygen-based oxidation enzymes. Instead, the aromatic rings arise through cyclization reactions of the polyketide chain. While oxygen atoms may appear in the final structures (e.g., as ketone or hydroxyl groups), they are typically derived from precursor molecules rather than external oxidation processes. Previous phylogenetic analyses suggested that nonoxidative T2PKSs diverged from other T2PKSs early during the evolutionary process because of their distinct protein sequences. However, our dimensionality reduction analysis of T2PKS PLM embeddings (Figure 1b) revealed that nonoxidative T2PKSs do not cluster with the ancestral sequences predicted by the MAPPE algorithm. Instead, these sequences are embedded within the KS cluster, labelled the CLF_4 cluster. The sequence IDs of KS and CLF for non-oxidative type II polyketides are documented in Supplementary Table S5 [7].

To further investigate the unexpected affinity between the CLF_4 and KS clusters and to elucidate the discrepancies between Cluster A and the previously published ancestral sequence predictions, we employed the DeepFRI utility to conduct functional predictions on the 166 KS-CLF sequence pairs (Supplementary Table S6) [20]. This deep learning-based functional annotation method promises to provide new insights into the functional evolution and clustering patterns of such proteins.

**FIGURE 3.**
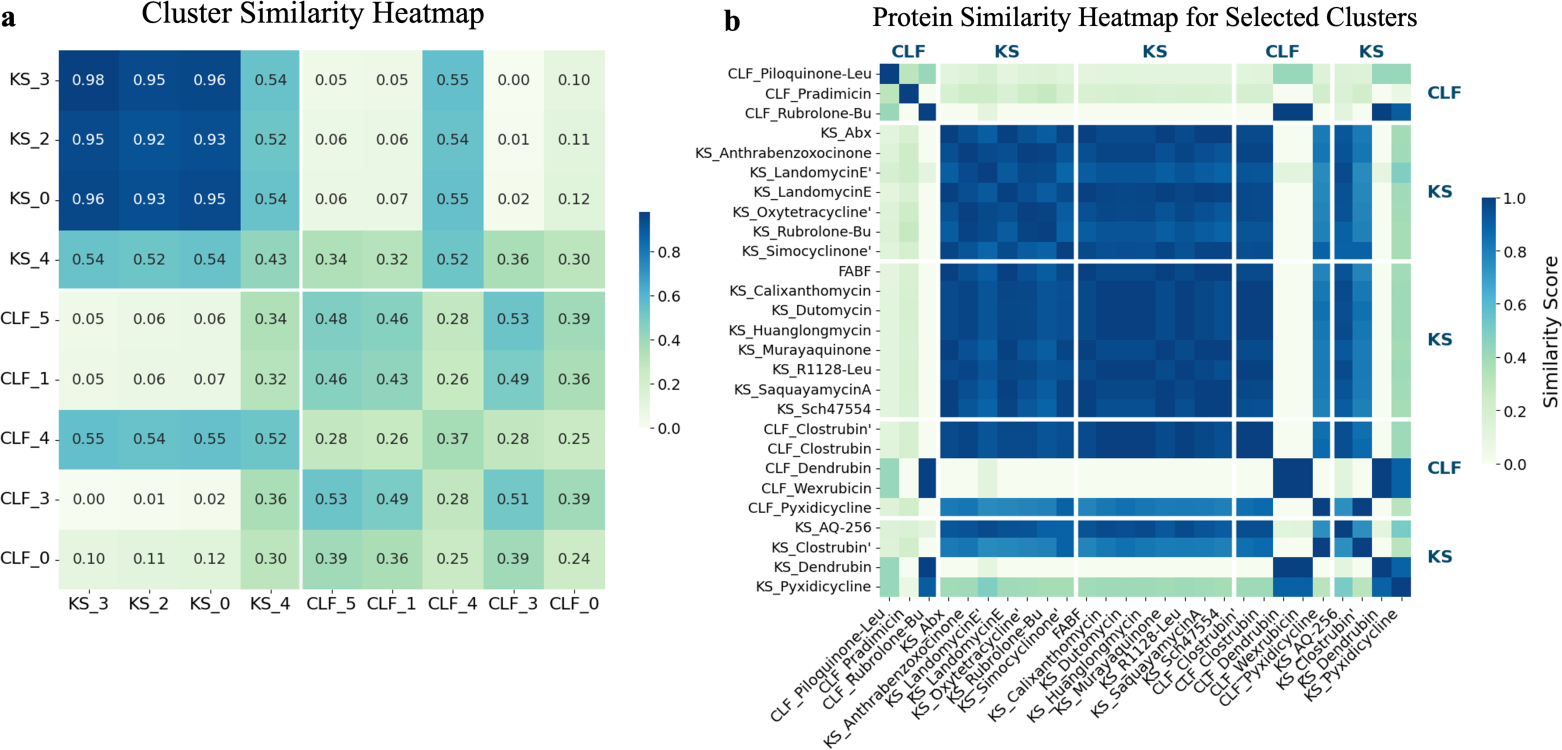
Function prediction and similarity analysis results obtained for 166 KS/CLF pairs by DeepFRI. Functional similarity is based on the cosine similarity of the vectorized GO classifications and probabilities. Higher similarity values indicate greater functional resemblance between proteins. Heatmap of the pairwise functional similarities among all proteins. Proteins were clustered based on the results shown in Figure 2b. The similarity scores were calculated via the cosine similarity between the one-hot encoded vectors of the DeepFRI-predicted GO terms and their probabilities. (b) Detailed similarity heatmaps for the proteins within Cluster A (CLF_4, KS_0, and KS_4) and the nonoxidative CLF_4 and KS_4 groups. The colour scale ranges from light green to deep blue, representing low to high functional similarity. This figure was created using BioRender.

Based on the cluster representations and intergroup evolutionary pathways predicted by edge bundling as shown in Figure 2c, CLF_4 appears to have evolved through a trajectory from CLF_0 to KS_0/KS_2/KS_3 and finally to CLF_4. Notably, a group of nodes embedded within the largest KS_2 cluster contained in the two-dimensional reduced representation did not cluster with KS_2. Instead, these nodes formed a distinct KS_4 cluster, which corresponds to the KS components of nonoxidative T2PKSs.

Figure 3 presents the functional predictions produced for the KS and CLF protein sequences through the DeepFRI analysis. As proteins often exhibit multiple functions, DeepFRI outputs GO categories with associated probabilities (0-1) for each protein. We represented each protein as a GO-term-based functional vector (one-hot), in which the dimensionality corresponds to all relevant GO categories and each dimension stores the probability (0-1) assigned by DeepFRI to that GO term for the given protein (Supplementary Table S7) [21].

The functional similarity values between the proteins were then quantified via the cosine similarity values between their respective functional vectors. We conducted pairwise similarity comparisons across all the KS and CLF clusters, with the results illustrated in Figure 3a. Detailed functional similarity matrix between proteins is listed in Supplementary Table S8 and Supplementary Figure S1. The KSs exhibited high functional similarity, indicating a conserved chain elongation function that plays a fundamental and crucial role in biological processes. In contrast, the CLFs demonstrated greater functional diversity, as reflected in lower pairwise functional similarities, which corresponds to their role in controlling chain lengths for achieving product diversity via adaptation to challenging environments.

Two CLF groups, CLF_0 and CLF_4, exhibited notably distinct similarity patterns in comparison with those of other CLF clusters. CLF_0, which is located in Cluster A, and CLF_4, which is associated with nonoxidative T2PKSs, displayed markedly greater functional similarity to the KS groups than the other CLF clusters did. Specifically, CLF_0 maintained high similarity with the other CLF groups while also being significant similar to the KS groups. CLF_4, however, is more similar to the KS groups than to the CLF groups. These functional prediction results provide an explanation for the root position of CLF_0 in the MAPPE network. Furthermore, they elucidate the unexpected distribution of the nonoxidative CLFs among KS nodes.

KS_4 represents an outlier within the KS group. As previously mentioned, it was distributed among the KS nodes but formed a distinct cluster during the clustering analysis. Its functional similarity to other KS clusters was less pronounced; instead, it showed comparable similarity to CLF_4 (approximately 0.5 for both the KS and CLF comparisons). Figure 3b presents a heatmap of the pairwise functional similarities among all proteins within the CLF_0, KS_0, KS_3, CLF_4, and KS_4 clusters. Notably, the CLF domains of clostrubin and pyxidicycline in CLF_4 closely resemble those of KSs. Conversely, the KS domains of these two compounds, along with Dendrubin in KS_4, display functions that are similar to those of CLFs (More information regarding the comparisons between other clusters, see Supplementary Figures S2 and S3). This functional overlap suggests that the KS and CLF domains in nonoxidative PKSs may not have fully differentiated, potentially retaining the characteristics of their early evolutionary states.

Our findings support the theory of a common origin for KSs and CLFs, which was followed by functional divergence to achieve product diversity. The DeepFRI output indicates that the primary function of a KS is GO:0016746: “transferase activity, transferring acyl groups.” [22] CLFs with KS-like functions also exhibit acetyltransferase activity, suggesting that in their early stages, both KSs and CLFs served to generate T2PK chains. To adapt to diverse environments, CLFs subsequently evolved various novel functions, facilitating the production of new T2PKSs and thus achieving product diversity.

This functional analysis offers critical insights into the evolutionary relationships and functional diversification trends of T2PKS components, reconciling the phylogenetic observations determined in the context of structural analogy with predicted protein functions.

#### 4. Co-evolution of KSs and CLFs

MAAPE offers a novel approach for detecting coevolutionary patterns. Figure 2a reveals a symmetric relationship between the KS and CLF groups. We assessed the ratio of the distances in the embedding space between each KS and CLF pair (originating from the same biosynthesis-related gene cluster) relative to point A, which represents the ancestral node, and defined it as the centroid of the embeddings in Cluster A (Figure 2d, Supplementary Table S9) [23]. The mean ratio was calculated to be 1.0082, which is very close to 1.0, suggesting that since diverging from their common ancestor, most KS and CLF pairs have evolved with nearly identical evolutionary distances and experienced similar selection pressures. Only six node pairs exhibited ratio differences greater than 0.2; in five of these cases, the KS distances were longer than the CLF distances, indicating differential selection pressures within these specific gene clusters. The overall symmetry and consistent distance ratios strongly support the hypothesis of coevolution between the KS and CLF genes within these biosynthetic pathways. From the viewpoint of biological interpretation, our analysis suggests that the divergence of KS-CLF clusters is likely shaped by multiple selective pressures. First, different clusters are associated with products exhibiting distinct bioactivities, implying chemical-mediated competition as a major driving force. Second, host organisms occupy diverse ecological niches, which may impose environment-specific constraints on secondary metabolism. Third, variations in scaffold complexity and tailoring reactions reflect trade-offs between metabolic cost and adaptive benefit. Together, these factors may contribute to the observed evolutionary trajectories of aromatic polyketide synthases.

**FIGURE 4.**
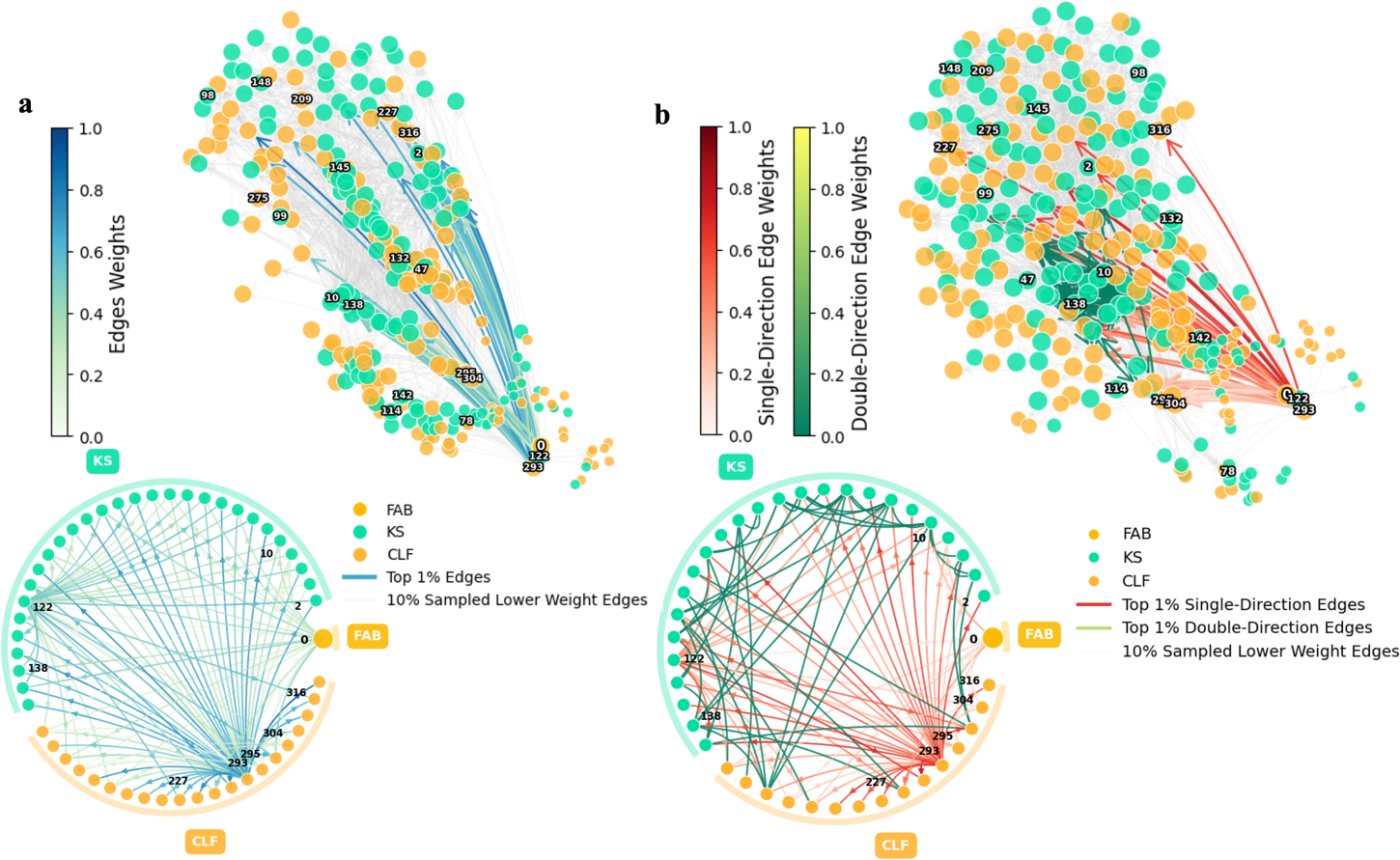
Visualization of the KS-CLF co-occurrence matrix. Only edges with weights in the top 1% are shown, 15% sampled background edges (gray) provide network context while highlighting the most significant connections. The nodes were categorized by colour. The similarities between nodes can be inferred from the edges connecting them. Designated directions are marked by arrows, and the node sizes reflect their degrees, with larger nodes indicating more intensive connections and greater importance levels in the network. Only Cluster-A labels are shown in the figure. Lower panels show chord diagram representations of the same top 1% highest-weighted connections accordingly. (a) This graph was derived by calculating the weights of the edges between pairs of nodes in both the forward and reverse directions, retaining only the direction with the larger weight. Considering that weights also represent similarity values, the sum of the bidirectional weights was used when mapping edge colours, and the edge weights are represented by colours ranging from light to deep blue, indicating higher similarity and more extensive genomic exchange signals. This suggests that these nodes have undergone frequent recombination or HGT events. (b) Uni- or bidirectional edge visualization. When the bidirectional edge weights were calculated, the direction of the edge with the larger weight was retained only if the difference exceeded 50%. If the difference was less than 50%, it was recorded as a unidirectional edge. Unidirectional and bidirectional edges were mapped via red and green bars, respectively, demonstrating how unidirectional and bidirectional connections are spatially segregated between different cluster regions, The MAAPE algorithm identifies nodes 0, 122 (KS_0), and 293 (CLF_0) as Cluster A members positioned at the root nodes of the network, which is consistent with their predicted ancestral roles in the weight matrix trajectory. This figure was created using BioRender.

#### 5. HGT is a main driving force of the KS/CLF polymorphism

We further visualized the co-occurrence matrix to assess the extent to which the MAAPE algorithm could capture evolutionary information. The results are displayed in Figure 4. In graph (a), directional arrows were assigned based on the higher edge weights between the pairs of nodes with existing connections, allowing the unidirectional arrows to directly represent the inferred evolutionary order. In graph (b), we compared the weights of both directions between node pairs. For edges with a weight difference exceeding 50%, a unidirectional arrow points in the direction of the higher weight; for differences below 50%, bidirectional arrows are used. The colour mappings of the edges in both graphs correspond to the total weight in both directions, whereas the node sizes reflect the numbers of connections, indicating the importance and centrality of nodes within the network structure. The sequence IDs within Cluster A are highlighted in both figures.

The nodes located in the lower-right corners of both graphs correspond to three nodes from Cluster A, notably including the FabF enzyme. This aligns with the current consensus that early forms of T2PKSs likely diverged from Fab ancestors and thus retained more Fab-like features, as reflected in both the evolutionary order inferred from MAAPE and the network structure. The outwardly expanding trends of these nodes, indicated by their connecting arrows, suggests that they emerged earlier in the evolutionary timeline and served as foundational points for the subsequent diversification process. Notably, the remaining members of Cluster A are characterized by larger node sizes, highlighting their significance within the network. The KS_0 and CLF_0 groups, which were predicted to occupy earlier evolutionary positions, are represented by particularly large nodes. This finding is consistent with their hypothesized ancestral statuses and key roles in the evolutionary process.

Interestingly, while KS_3 is also part of Cluster A and is located close to KS_0 and CLF_0, it displays a different pattern. The nodes representing KS_3 are more dispersed and smaller than their cluster counterparts. This distinctive characteristic suggests that KS_3 may represent an early diversification attempt in the evolutionary history of KSs and CLFs. Although its structural features were preserved, it did not result in significant expansion akin to that of its counterparts.

In Figure 4b, evolutionary relationships are visualized with directional arrows and colour-coded weights. Unidirectional arrows are represented by a gradient of red, ranging from light to dark, whereas bidirectional arrows are shown in shades of green. The darker red unidirectional arrows, which are predominantly located near the ancestry nodes, suggest that KSs and CLFs may have originally evolved from a few ancestral sequences with clear directional trends. This is supported by the high weights assigned to HGT and other gene module transfer events in MAAPE, indicating that such events were more prevalent during the early stages of evolution.

As we move towards the smaller nodes and the outer boundaries of the network, the arrow colours become lighter, signifying less impactful or more recent evolutionary events. In these regions, an increase in the number of bidirectional green arrows suggests more frequent recombination or point mutation events. This evolutionary strategy, which is particularly evident in the early stages with darker red arrows, likely facilitated the pioneering of novel natural products with new functions or carbon skeletons. The transition from predominantly unidirectional dark red arrows to a mix of lighter red and green arrows reflects a shift from bold evolutionary changes to more nuanced modifications and bidirectional genetic exchanges. This pattern illustrates the initial exploration of new functional spaces, followed by fine-tuning and diversification within the established structural frameworks.

A notable observation derived from the network is the presence of CLF_4 within the KS cluster, suggesting that HGT may have conferred KS-like properties and functions upon these genes despite their distant sequence similarities. The ESM-2 model appears to have effectively captured these transferred features. In Figure 2e, the edges near these nodes are light yellow, indicating high colour mapping weights and correlating with strong HGT signals. Specifically, light yellow edges connect the upper-left corner of the large CLF cluster with Cluster A and CLF_4, explaining the acquisition of KS-like properties. This discovery underscores the significant role of HGT in the PKS evolution process, leading to similar substrate or product specificities. Further investigations into these nodes could provide insights into the origins of novel PKS functions. Analysing the species in which these HGT events occurred, along with their changing environmental conditions, could illuminate the driving forces behind these evolutionary processes.

## Discussions

The evolutionary process of bacterial PKSs has been extensively studied over the past few decades, but this has primarily been done through phylogenetic bioinformatic approaches. In this work, we applied different strategies to reconstitute the evolutionary relationships of 2 structurally homologous proteins: the KS and CLF domains derived from bacterial T2PKSs. We identified a potential ancestral group (Cluster A) containing both KS and CLF sequences, supporting the hypothesis of a common origin. Following their initial divergence, we observed distinct evolutionary trajectories for KSs and CLFs, with evidence of coevolution within biosynthesis-related gene clusters. The calculated weight matrix guided the subsequent detection and analysis of the critical sequence modules, potentially offering new perspectives for the design and synthesis of novel natural products. Through the prediction of protein functions via DeepFRI, we found that CLFs predicted with earlier evolutionary positions in our analysis have functions similar to those of KSs, thus explaining the unexpected clustering results of nonoxidative CLFs and KSs. These findings verify that structurally homologous proteins have certain functional overlaps but can evolve into highly divergent sequences with conserved structures during adaptations to different environments. The significant role of HGT was detected in the early evolutionary stages, shaping the diversity of KSs and CLFs, which also led to potential instances of functional convergence between nonoxidative CLFs and KSs, suggesting complex evolutionary dynamics [23].

The theory of evolution viewed through the assembly of sequence modules suggests that biological entities evolve convergently across different organisms by exchanging useful sequence fragments rather than generating essential modules from scratch [24]. This process conserves energy and accelerates environmental adaptations, with the repeated occurrence of specific fragments indicating conserved or crucial functional roles. To leverage the power of PLMs and advanced computational techniques to explore whether different-sized slices of embedded protein sequences carry meaningful evolutionary signals, we have recently developed MAPPE, a novel module-based evolutionary recognition and directional computation algorithm, aimed at (1) predicting the existence of ancestral or evolutionary root systems within a network of evolutionary relationships; (2) identifying the gene flow direction during an evolutionary process, where basic modules are assembled or transferred to form larger modules with more complex functions; (3) locating the crucial sequence modules that play pivotal roles in key bifurcation events by segmenting the embedded sequences and identifying the critical sequence segments; and (4) inferring the functional and evolutionary links between proteins and the driving forces behind the instances that shape these trajectories.

Compared with the traditional phylogenetic analysis process, the MAAPE algorithm, as established in this work, offers greater spatial resolutions for the relationships between different proteins. This is because the traditional methods require the integration of information from multiple independent analyses, whereas the generated network graphs provide insights into coevolutionary processes and the magnitudes of selective pressures (among other factors) and offer comprehensive all-in-one network visualizations. We believe that further in-depth analyses of these networks will yield even richer information.

Finally, and importantly, the connection between the computational results and their biological interpretation cannot be overlooked. As discussed in the results section, from a biological perspective, our analysis indicates that the divergence of KS and CLF clusters is influenced by multiple selective pressures, including chemical competition among microorganisms driving functional diversification, environmental constraints on secondary metabolism imposed by hosts occupying different ecological niches, and trade-offs between scaffold complexity and tailoring reactions that shape enzyme evolution to balance metabolic efficiency with chemical versatility.

Taken together, these findings suggest that the evolutionary trajectories of aromatic polyketide synthases are shaped by a combination of chemical competition, ecological pressures, metabolic trade-offs, structural and functional constraints, and horizontal gene exchange. The integration of sequence, structural, and functional data through PLM embeddings provides a more nuanced view of how KS and CLF domains have diversified to generate the remarkable chemical diversity observed in bacterial secondary metabolites.

## Conclusions

To conclude, examining the evolutionary connections between KSs and CLFs provides a unique window into the inner workings of bacterial chemistry. Understanding the shared history, functions, and intricacies of KS-CLF interactions not only deepens our comprehension of microbial life but also provides immense potential for rationally designing enzymes and manipulating PKS assembly lines, ultimately benefiting a spectrum of fields ranging from medicine to biotechnology.

## Methods

### Data collection and phylogenetic analysis

Phylogenetic analysis was performed using a curated dataset of 166 pairs of bacterial KS-CLF proteins with known polyketide product structures, along with a *Staphylococcus aureus* FABF sequence as an outgroup. Multiple sequence alignment was conducted using CLUSTALW [25]. The phylogenetic tree was constructed using IQ-TREE with the LG+F+R8 substitution model, which was selected as the best-fit model based on the Bayesian information criterion [26]. Tree visualization and aesthetic refinement were performed using the Interactive Tree of Life (iTOL) web-based tool [27].

### Structure analysis

The structures of KS and CLF proteins were predicted using the AlphaFold Protein Structure Database server [16]. Structural comparisons were performed using PyMOL [28]. The structures were first prepared by removing heteroatoms and water molecules, followed by structure alignment. Root-mean-square deviation (RMSD) values were calculated based on Cα atomic positions.

### Protein language model embedding

To analyse protein sequence patterns, we employed the ESM-2 protein language model, which implements a Transformer-based deep learning architecture [10]. We utilized the pre-trained ESM2_t36_3B_UR50D model from Meta AI, featuring 36 transformer layers and approximately 3 billion parameters. The model processes protein sequences through a tokenization step, followed by contextual encoding across multiple attention layers. For each protein sequence in our dataset, we extracted the representations from the final hidden layer (dimension 2560) as sequence-specific feature vectors. These embeddings were subsequently normalized using L2 normalization to ensure consistent comparison across different sequences. Embeddings were subsequently reduced to three dimensions using Uniform Manifold Approximation and Projection (UMAP) algorithm [29].

To address the computational load posed by the high-dimensional embedding vectors (2560 dimensions), we employed Principal Component Analysis (PCA) for initial dimensionality reduction [30]. Analysis of the cumulative explained variance revealed that reducing the embeddings to 220 principal components preserved more than 60% of the original data variance, providing an optimal balance between computational efficiency and information retention.

### MAAPE analysis

We developed and implemented MAAPE methodology to extract evolutionary relationships from protein language model embeddings while overcoming traditional sequence alignment limitations. The analysis framework consists of two primary components: a similarity network construction and a directional assembly pattern detection.

For the similarity network, we constructed a k-nearest neighbor (KNN) graph based on Euclidean distances between embedded vectors [31]. The network topology was computed using the IndexFlatL2 algorithm implemented in Faiss library, which performs exact L2 distance calculations to ensure precise similarity measurements [32]. This approach captures diverse evolutionary relationships including functional divergence, structural modifications, and sequence variations.

To analyse assembly patterns, we developed a sliding window approach where protein embeddings were systematically segmented into variable-length fragments, starting from smaller sub-vectors and progressively moving to larger ones, we computed pairwise Euclidean distances between sub-vectors to identify similarity and containment relationships. A co-occurrence matrix was constructed based on these relationships, the matrix elements represent both the similarity between embedding segments and their hierarchical relationships. Assembly directionality was established by tracking the relationships between shorter segments and their containing longer fragments, reflecting the progressive organization of evolutionary features. This dual-component analysis enables the detection of both global sequence relationships and local evolutionary patterns, providing insights into protein evolution that may not be apparent through conventional sequence alignment methods.

### Functional prediction

Protein function prediction was performed using DeepFRI, a deep learning-based functional annotation tool [20]. For each KS and CLF protein sequence, DeepFRI generated predictions across multiple Gene Ontology (GO) categories with associated probability scores, we only focus on their molecular function output in this work [21]. To enable quantitative comparison of functional similarities between proteins, we developed a vectorization approach where the predicted GO terms and their corresponding probability scores were transformed into fixed-length vectors using one-hot encoding. This encoding process created a standardized representation for each protein’s functional profile, where each dimension corresponds to a specific GO term and its prediction confidence.

Functional similarities between protein pairs were quantified using cosine similarities, where higher values indicate greater functional closeness between proteins. The pairwise similarities were visualized using clustered heatmaps, with clustering based on the MAAPE output relationships.

## Data availability

The authors declare that the data, materials and code supporting the findings reported in this study are publicly available. The MAAPE is available at GitHub repository: https://github.com/Qinlab502/MAAPE, and archived at Zenodo (https://doi.org/10.5281/zenodo.18631852).

## Author contributions

Zhiwei Qin designed and supervised the research. Xiaoyu Wang performed the bioinformatic and established the algorithm. Qiandi Gao and Liangjun Ge performed the bioinformatic and chemical analysis. All authors analysed and discussed the data. Zhiwei Qin and Xiaoyu Wang wrote the manuscript and all authors edited.

## Acknowledgements

The authors would like to thank the Interdisciplinary Intelligence Super Computer Center, Beijing Normal University, for High Performance Computing for access to computational resources. We acknowledge BioRender for providing the illustration platform used in this study.

## Declarations

### Ethics approval and consent to participate

Not applicable.

### Consent for publication

Not applicable.

