## supplementary information for "The evolutionary processes of bacterial aromatic polyketide ketosynthases"

ELECTRONIC SUPPLEMENTARY INFORMATION

**Table of contents**

1. ESI Tables

2. ESI Figures

### 1. ESI Tables

**Supplementary Table S1.** Index and strain information for all 166 T2 PKS KS-CLF pairs in this study. No. 1-166 is assigned to KSs, No. 167-332 is assigned to their corresponding CLFs.

| Index | T2PK products name | Building block number | Strain | Accession |
|---|---|---|---|---|
| 0 | FABF | | *Staphylococcus aureus* | BAD72839.1 |
| 1 | ksa_A-74528-Hex_13 | 13 | *Streptomyces sp. SANK 61196* | ADG86315.1 |
| 2 | ksa_Abx_13 | 13 | *Streptomyces sp.* | AUI41025.1 |
| 3 | ksa_+ABXA-BE-24566B_13 | 13 | *Streptomyces sp.* | KU243130.1 |
| 4 | ksa_Accramycin_13 | 13 | *Streptomyces sp.* | QKO28696.1 |
| 5 | ksa_Aclacinomycin'-Pr_10 | 10 | *Streptomyces galilaeus* | AAF70106.1 |
| 6 | ksa_Aclacinomycin-Pr_10 | 10 | *Streptomyces galilaeus* | BAB72045.1 |
| 7 | ksa_Actinorhodin_8 | 8 | *Streptomyces coelicolor A3(2)* | CAC44200.1 |
| 8 | ksa_Allocyclinone_10 | 10 | *Actinoallomurus sp. ID145698* | AMX23327.1 |
| 9 | ksa_Alnumycin-Bu_8 | 8 | *Streptomyces sp. CM020* | ACI88861.1 |
| 10 | ksa_Anthrabenzoxocinone_13 | 13 | *Actinobacteria bacterium* | AUO15560.1 |
| 11 | ksa_AQ-256_8 | 8 | *Photorhabdus laumondii subsp. laumondii TTO1* | CAE16563.1 |
| 12 | ksa_Aranciamycin_10 | 10 | *Streptomyces echinatus* | ABL09959.1 |
| 13 | ksa_Arenimycin'_12 | 12 | *uncultured bacterium BAC-AB1442/1414/561* | AIW63010.1 |
| 14 | ksa_Arenimycin_12 | 12 | *Salinispora arenicola* | WP_029025958.1 |
| 15 | ksa_Arimetamycin_10 | 10 | *uncultured bacterium* | AHA81977.1 |
| 16 | ksa_Arixanthomycin_13 | 13 | *uncultured bacterium* | AHX24701.1 |
| 17 | ksa_Auricin_10 | 10 | *Streptomyces lavendulae subsp. lavendulae* | AAX57191.1 |

| | | | | |
|---|---|---|---|---|
| 18 | ksa_Azicemicin-aziridine_10 | 10 | *Kibdelosporangium sp. MJ126-NF4* | ADB02843.1 |
| 19 | ksa_Baikalomycin_10 | 10 | *Streptomyces sp. IB201691-2A2* | WP_143644093.1 |
| 20 | ksa_BE-7585A_10 | 10 | *Amycolatopsis orientalis subsp. vinearia* | ADI71443.1 |
| 21 | ksa_Benastatin-Hex_12 | 12 | *Streptomyces sp. A2991200* | CAM58798.1 |
| 22 | ksa_Bipentaromycin_10 | 10 | *Streptomyces sp. NRRL F-6131* | WP_030301373.1 |
| 23 | ksa_Brasiliquinone-Pr_10 | 10 | *Nocardia brasiliensis* | WP_146154178.1 |
| 24 | ksa_Calixanthomycin_12 | 12 | *uncultured bacterium* | AJD20010.1 |
| 25 | ksa_Cervimycin-malonoamyl_9 | 9 | *Streptomyces sp. CS113* | OWA01596.1 |
| 26 | ksa_Chartreusin'_10 | 10 | *Streptomyces chartreusis* | AXS67802.1 |
| 27 | ksa_Chartreusin_10 | 10 | *Streptomyces chartreusis* | CAH10161.1 |
| 28 | ksa_Chattamycin_10 | 10 | *Streptomyces lydicus* | WP_046929079.1 |
| 29 | ksa_Chelocardin_10 | 10 | *Amycolatopsis sulphurea* | AHD25926.1 |
| 30 | ksa_Chlortetracycline_10 | 10 | *Kitasatospora aureofaciens* | AEI98666.1 |
| 31 | ksa_Chromomycin_10 | 10 | *Streptomyces griseus subsp. griseus* | CAE17527.1 |
| 32 | ksa_Chrysomycin-Pr_10 | 10 | *Streptomyces albaduncus* | CBH32088.1 |
| 33 | ksa_Cinerubin-Pr_10 | 10 | *Streptomyces sp. SPB074* | EDY42532.1 |
| 34 | ksa_Clostrubin'_X | 10 | *Clostridium puniceum* | WP_077846908.1 |
| 35 | ksa_Clostrubin_X | 10 | *Clostridium beijerinckii* | WP_077837156.1 |
| 36 | ksa_Collinone_13 | 13 | *Streptomyces collinus* | AAG26879.1 |
| 37 | ksa_CosmomycinC-Pr_10 | 10 | *Streptomyces sp. CNT302* | WP_017947381.1 |
| 38 | ksa_Cosmomycin-Pr_10 | 10 | *Streptomyces olindensis* | ABC00726.1 |
| 39 | ksa_Cur_12 | 12 | *Streptomyces cyaneus* | CAA44380.1 |
| 40 | ksa_Cysgriseusin_10 | 10 | *Streptomyces cyaneofuscatus* | WP_030562167.1 |

| | | | | |
|---|---|---|---|---|
| 41 | ksa_Cytorhodin-Pr_10 | 10 | *Streptomyces sp. SCSIO 1666* | ATJ00755.1 |
| 42 | ksa_Dactylocycline_10 | 10 | *Dactylosporangiu m sp. SC14051* | AFU65894.1 |
| 43 | ksa_Daunorubicin-Pr_10 | 10 | *Streptomyces sp.* | AAA87618.1 |
| 44 | ksa_Dendrubin_9 | 9 | *Dendrosporobacte r quercicolus* | WP_092069528 .1 |
| 45 | ksa_Doxorubicin'-Pr_10 | 10 | *Streptomyces peucetius* | WP_100109102 .1 |
| 46 | ksa_Doxorubicin-Pr_10 | 10 | *Streptomyces peucetius* | AAA65206.1 |
| 47 | ksa_Dutomycin_10 | 10 | *Streptomyces minoensis* | AKD43514.1 |
| 48 | ksa_Elloramycin_10 | 10 | *Streptomyces olivaceus* | CAP12600.1 |
| 49 | ksa_Enduracyclinone_13 | 13 | *Nonomuraea sp.* | AYP71359.1 |
| 50 | ksa_Enterocin-Bn_8 | 8 | *Streptomyces maritimus* | AAF81728.1 |
| 51 | ksa_Erdacin_8 | 8 | *uncultured soil bacterium V167* | ACX83617.1 |
| 52 | ksa_Fasamycin_13 | 13 | *N/A* | AEM44268.1 |
| 53 | ksa_FD-594-Bu_13 | 13 | *Streptomyces sp. TA-0256* | BAJ52681.1 |
| 54 | ksa_Fluostatin_10 | 10 | *uncultured bacterium BAC AB649/1850* | AEE65468.1 |
| 55 | ksa_FluostatinM_10 | 10 | *Micromonospora rosaria* | ALJ99855.1 |
| 56 | ksa_FluostatinMQ_10 | 10 | *Streptomyces albus* | WP_040253443 .1 |
| 57 | ksa_Fogacin_8 | 8 | *Actinoplanes missouriensis 431* | BAL90284.1 |
| 58 | ksa_FogacinC-HCS_8 | 8 | *Actinoplanes missouriensis 431* | BAL90263.1 |
| 59 | ksa_Formicamycin_13 | 13 | *Streptomyces sp. KY5* | AQP25565.1 |
| 60 | ksa_Frankiamicin_12 | 12 | *Frankia sp. EAN1pec* | ABW11825.1 |
| 61 | ksa_FredericamycinC_13 | 13 | *Streptomyces albus subsp. chlorinus* | NSC21571.1 |
| 62 | ksa_Fredericamycin-Hex_13unknown | 13 | *Streptomyces griseus* | AAQ08916.1 |
| 63 | ksa_Frenolicin-Bu_8unknown | 8 | *Streptomyces roseofulvus* | AAC18107.1 |
| 64 | ksa_Frigocyclinone_10unknown | 10 | *Streptomyces griseus* | QDG00826.1 |

| | | | | |
|---|---|---|---|---|
| 65 | ksa_Gaudimycin_10 | 10 | *Streptomyces sp. PGA64* | AAK57525.1 |
| 66 | ksa_Gilvocarcin-Pr_10 | 10 | *Streptomyces griseoflavus* | AAP69573.1 |
| 67 | ksa_Granaticin''_8unknown | 8 | *Streptomyces exfoliatus* | WP_137988251.1 |
| 68 | ksa_Granaticin'_8unknown | 8 | *Streptomyces violaceoruber* | CAA09653.1 |
| 69 | ksa_Granaticin_8unknown | 8 | *Streptomyces vietnamensis* | ADO32786.1 |
| 70 | ksa_Grincamycin_10 | 10 | *Streptomyces lusitanus* | AGO50610.1 |
| 71 | ksa_Griseorhodin_13 | 13 | *Streptomyces sp. JP95* | AAM33653.1 |
| 72 | ksa_Griseusin_10unknown | 10 | *Streptomyces griseus* | CAA54858.1 |
| 73 | ksa_Hatomarubigin_10unknown | 10 | *Streptomyces sp. 2238-SVT4* | BAJ07842.1 |
| 74 | ksa_Hedamycin-Hex_10 | 10 | *Streptomyces griseoruber* | AAP85362.1 |
| 75 | ksa_Heliquinomycin_13unknown | 13 | *Streptomyces piniterrae* | WP_136741436. |
| 76 | ksa_Hexaricin_13 | 13 | *Streptosporangium sp. FXJ7.131* | AMK51281.1 |
| 77 | ksa_Hiroshidine_8 | 8 | *Streptomyces hiroshimensis* | QBK46637.1 |
| 78 | ksa_Huanglongmycin_9unknown | 9 | *Streptomyces sp. CB09001* | AXL88816.1 |
| 79 | ksa_Hyaluromycin_13unknown | 13 | *Streptomyces hyaluromycini* | WP_089099014. |
| 80 | ksa_Hydroxyfujianmycin_10unknown | 10 | *Streptomyces alni* | WP_093715797. |
| 81 | ksa_IFQs_8 | 8 | *Streptomyces sp. CB01883* | APR73637.1 |
| 82 | ksa_Isatropolone-Bu_8unknown | 8 | *Streptomyces sp.* | ARG41911.1 |
| 83 | ksa_Isoindolinomycin-Gly_8unknown | 8 | *Streptomyces sp. SoC090715LN-16* | BBE36466.1 |
| 84 | ksa_Jadomycin_10unknown | 10 | *Streptomyces venezuelae ATCC 10712* | AAB36562.1 |
| 85 | ksa_JBIR-76_8 | 8 | *Streptomyces sp. RI-77* | BAU98041.1 |
| 86 | ksa_Julichrome'_9unknown | 9 | *Streptomyces sampsonii* | QNL10613.1 |
| 87 | ksa_Julichrome_9unknown | 9 | *Streptomyces afghaniensis* | WP_020275095. |
| 88 | ksa_Keyicin_10unknown | 10 | *Micromonospora sp. WMMB235* | OHX01516.1 |

| | | | | |
|---|---|---|---|---|
| 89 | ksa_Kiamycin_9unknown | 9 | *Streptomyces sp. W007* | EHM27505.1 |
| 90 | ksa_Kinamycin_10unknown | 10 | *Streptomyces murayamaensis* | AAO65346.1 |
| 91 | ksa_kinanthraquinone_9unknown | 9 | *Streptomyces sp. SN-593* | WP_202233564. |
| 92 | ksa_Komodoquinone'_10unknown | 10 | *Streptomyces erythrochromogenes* | WP_031147015. |
| 93 | ksa_Komodoquinone_10unknown | 10 | *Streptomyces sp. NRRL S-378* | WP_030957354.1 |
| 94 | ksa_Kosinostatin_10 | 10 | *Micromonospora sp. TP-A0468* | AFJ52674.1 |
| 95 | ksa_Lactonamycin-Gly_10 | 10 | *Streptomyces rishiriensis* | ABX71114.1 |
| 96 | ksa_LactonamycinZ-Gly_10 | 10 | *Streptomyces sanglieri* | ABX71148.1 |
| 97 | ksa_Landomycin_10unknown | 10 | *Streptomyces cyanogenus* | AAD13536.1 |
| 98 | ksa_LandomycinE'_10unknown | 10 | *uncultured bacterium* | AEM44218.1 |
| 99 | ksa_LandomycinE_10unknown | 10 | *Streptomyces globisporus* | AID46990.1 |
| 100 | ksa_LL-D49194_1-Ile_10 | 10 | *Streptomyces vinaceusdrappus* | QDQ37910.1 |
| 101 | ksa_Lomaiviticin''-Pr_10unknown | 10 | *Salinispora pacifica* | AHZ61892.1 |
| 102 | ksa_Lomaiviticin'-Pr_10unknown | 10 | *Salinispora pacifica* | AHF72794.1 |
| 103 | ksa_Lomaiviticin-Pr_10unknown | 10 | *Salinispora tropica CNB-440* | ABP54674.1 |
| 104 | ksa_Lugdunomycin_10unknown | 10 | *Streptomyces sp. QL37* | PPQ57489.1 |
| 105 | ksa_Lysolipin_12 | 12 | *Streptomyces tendae* | CAM34345.1 |
| 106 | ksa_Maduralactomycin_10unknown | 10 | *Actinomadura rubteroloni* | WP_103564581.1 |
| 107 | ksa_Mayamycin_10 | 10 | *Streptomyces sp.* | AVO00815.1 |
| 108 | ksa_Medermycin_8 | 8 | *Streptomyces sp. AM-7161* | BAC79044.1 |
| 109 | ksa_Mensacaricin_10unknown | 10 | *Streptomyces bottropensis* | AHL46686.1 |
| 110 | ksa_Metamycin-iPr_89unknown | 89 | *Rothia dentocariosa* | VEJ30651.1 |
| 111 | ksa_Metathramycin_10unknown | 10 | *uncultured bacterium* | QVQ68799.1 |
| 112 | ksa_Methyltetrangomycin_10unknown | 10 | *Streptomyces sp. Go-475* | WP_114256228.1 |

| | | | | |
|---|---|---|---|---|
| 113 | ksa_Mithramycin_10unknown | 10 | *Streptomyces argillaceus* | CAA61989.1 |
| 114 | ksa_Murayaquinone_10unknown | 10 | *Streptomyces griseoruber* | AQW35058.1 |
| 115 | ksa_Naphthocyclinone_8unknown | 8 | *Streptomyces arenae* | AAD20267.1 |
| 116 | ksa_Nenestatin-Pr_10unknown | 10 | *Micromonospora echinospora* | ARD70901.1 |
| 117 | ksa_Nivetetracyclate-Pr_10unknown | 10 | *Streptomyces sp. Ls2151* | AGZ78377.1 |
| 118 | ksa_Nocardiopsistin-iPr_10unknown | 10 | *Nocardiopsis sp.* | QOP59270.1 |
| 119 | ksa_Nogalamycin_10unknown | 10 | *Streptomyces nogalater* | CAA12017.1 |
| 120 | ksa_Oviedomycin'_10 | 10 | *Streptomyces antibioticus* | ARK36155.1 |
| 121 | ksa_Oviedomycin_10 | 10 | *Streptomyces antibioticus* | CAG14965.1 |
| 122 | ksa_Oxytetracycline'_10unknown | 10 | *Streptomyces rimosus subsp. rimosus ATCC 10970* | ELQ83269.1 |
| 123 | ksa_Oxytetracycline_10unknown | 10 | *Streptomyces rimosus* | AAZ78325.1 |
| 124 | ksa_Paramagnetoquinone_Xunknown | 10 | *Actinoallomurus sp.* | ASA49564.1 |
| 125 | ksa_PD116740_10unknown | 10 | *Streptomyces sp. WP 4669* | AAO65362.1 |
| 126 | ksa_Persiamycin_10unknown | 10 | *Streptomonospora sp. PA3* | WP_156000395.1 |
| 127 | ksa_Piloquinone-Leu_9 | 9 | *Streptomyces sp. FXJ8.102* | QFS19044.1 |
| 128 | ksa_Polyketomycin_10 | 10 | *Streptomyces diastatochromogenes* | ACN64834.1 |
| 129 | ksa_Pradimicin_12unknown | 12 | *Actinomadura hibisca* | ABM21747.1 |
| 130 | ksa_Pyxidicycline_9 | 9 | *Pyxidicoccus fallax* | AXM42927.1 |
| 131 | ksa_Qinimycin_8unknown | 8 | *Streptomyces roseifaciens* | WP_058047374.1 |
| 132 | ksa_R1128-Leu_8unknown | 8 | *Streptomyces sp. R1128* | AAG30189.1 |
| 133 | ksa_Ravidomycin-Pr_10unknown | 10 | *Streptomyces ravidus* | CBH32809.1 |
| 134 | ksa_Resistomycin_X | 10 | *Streptomyces resistomycificus* | CAE51174.1 |
| 135 | ksa_Rhodomycin-Pr_10unknown | 10 | *Streptomyces purpurascens* | WP_189725834.1 |

| | | | | |
|---|---|---|---|---|
| 136 | ksa_Rishirilide'-Leu_9unknown | 9 | *Streptomyces olivaceus* | AWM72904.1 |
| 137 | ksa_Rishirilide-Leu_9unknown | 9 | *Streptomyces bottropensis* | AHL46710.1 |
| 138 | ksa_Rubrolone-Bu_8unknown | 8 | *Streptomyces sp. KIB-H033* | AOZ61212.1 |
| 139 | ksa_Rubromycin_13 | 13 | *Streptomyces collinus* | WP_073806072.1 |
| 140 | ksa_Rubromycin'_13unknown | 13 | *unclassified Streptomyces* | AAG03067.1 |
| 141 | ksa_Saprolmycin_10unknown | 10 | *Streptomyces sp. TK08046* | BAV16999.1 |
| 142 | ksa_SaquayamycinA_10 | 10 | *Streptomyces sp.* | ARO44655.1 |
| 143 | ksa_SaquayamycinZ_10 | 10 | *Micromonospora sp. Tu 6368* | ACP19353.1 |
| 144 | ksa_Sch_12unknown | 12 | *Streptomyces halstedii* | AAA02833.1 |
| 145 | ksa_Sch47554_10unknown | 10 | *Streptomyces sp. SCC 2136* | CAH10117.1 |
| 146 | ksa_Setomimycin_9unknown | 9 | *Streptomyces aurantiacus* | WP_037658548.1 |
| 147 | ksa_SF2575_10 | 10 | *Streptomyces sp. SF2575* | ADE34518.1 |
| 148 | ksa_Simocyclinone'_10unknown | 10 | *Streptomyces sp. NRRL B-24484* | WP_052390556.1 |
| 149 | ksa_Simocyclinone_10unknown | 10 | *Streptomyces antibioticus* | AAK06784.1 |
| 150 | ksa_Steffimycin_10unknown | 10 | *Streptomyces steffisburgensis* | CAJ42320.1 |
| 151 | ksa_Streptoketide-Pr_10unknown | 10 | *Streptomyces sp. Tue6314* | QED90637.1 |
| 152 | ksa_Tetarimycin_10 | 10 | *uncultured bacterium* | AFY23044.1 |
| 153 | ksa_Tetracenomycin_10 | 10 | *Streptomyces glaucescens* | AAA67515.1 |
| 154 | ksa_Thioangucycline_10 | 10 | *Streptomyces sp. CB00072* | WP_073867016.1 |
| 155 | ksa_TLN-05220-Ile_13 | 13 | *Micromonospora echinospora subsp. challisensis* | ADB23391.1 |
| 156 | ksa_Trioxacarcin-Ile_10 | 10 | *Streptomyces bottropensis* | AKT74262.1 |
| 157 | ksa_Turbinmicin_13 | 13 | *Micromonospora sp. WMMC415* | WP_155333336.1 |
| 158 | ksa_Urdamycin_10 | 10 | *Streptomyces fradiae* | CAA60569.1 |
| 159 | ksa_UT-X26F129_10 | 10 | *Streptomyces aureus* | AYU66233.1 |

| | | | | |
|---|---|---|---|---|
| 160 | ksa_Warkmycin_10 | 10 | *Streptomyces sp. CS057* | OWA25245.1 |
| 161 | ksa_Wexrubicin_X | 10 | *Eubacteriales* | WP_025578803.1 |
| 162 | ksa_WhiE_12 | 12 | *Streptomyces coelicolor* | CAA39408.1 |
| 163 | ksa_WS79089D_13 | 13 | *Streptosporangium roseum DSM 43021* | WP_2890796.1 |
| 164 | ksa_X26_10 | 10 | *uncultured bacterium* | AEM44309.1 |
| 165 | ksa_Xantholipin_13 | 13 | *Streptomyces flavogriseus* | ADE22315.1 |
| 166 | ksa_ZHMpentangular_13 | 13 | *Streptomyces viridochromogenes* | WP_003993028.1 |
| 167 | ksb_A-74528-Hex_13 | 13 | *Streptomyces sp. SANK 61196* | ADG86316.1 |
| 168 | ksb_Abx_13 | 13 | *Streptomyces sp.* | AUI41026.1 |
| 169 | ksb_+ABXA-BE-24566B_13 | 13 | *Streptomyces sp.* | KU243130.1 |
| 170 | ksb_Accramycin_13 | 13 | *Streptomyces sp.* | QKO28695.1 |
| 171 | ksb_Aclacinomycin'-Pr_10 | 10 | *Streptomyces galilaeus* | AAF70107.1 |
| 172 | ksb_Aclacinomycin-Pr_10 | 10 | *Streptomyces galilaeus* | BAB72046.1 |
| 173 | ksb_Actinorhodin_8 | 8 | *Streptomyces coelicolor A3(2)* | CAC44201.1 |
| 174 | ksb_Allocyclinone_10 | 10 | *Actinoallomurus sp. ID145698* | AMX23328.1 |
| 175 | ksb_Alnumycin-Bu_8 | 8 | *Streptomyces sp. CM020* | ACI88862.1 |
| 176 | ksb_Anthrabenzoxocinone_13 | 13 | *Actinobacteria bacterium* | AUO15561.1 |
| 177 | ksb_AQ-256_8 | 8 | *Photorhabdus laumondii subsp. laumondii TTO1* | CAE16562.1 |
| 178 | ksb_Aranciamycin_10 | 10 | *Streptomyces echinatus* | ABL09958.1 |
| 179 | ksb_Arenimycin'_12 | 12 | *uncultured bacterium BAC-AB1442/1414/561* | AIW63011.1 |
| 180 | ksb_Arenimycin_12 | 12 | *Salinispora arenicola* | WP_029025959.1 |
| 181 | ksb_Arimetamycin_10 | 10 | *uncultured bacterium* | AHA81978.1 |
| 182 | ksb_Arixanthomycin_13 | 13 | *uncultured bacterium* | AHX24702.1 |

| | | | | |
|---|---|---|---|---|
| 183 | ksb_Auricin_10 | 10 | *Streptomyces lavendulae subsp. lavendulae* | AAX57192.2 |
| 184 | ksb_Azicemicin-aziridine_10 | 10 | *Kibdelosporangium sp. MJ126-NF4* | ADB02844.1 |
| 185 | ksb_Baikalomycin_10 | 10 | *Streptomyces sp. IB201691-2A2* | WP_143644094.1 |
| 186 | ksb_BE-7585A_10 | 10 | *Amycolatopsis orientalis subsp. vinearia* | ADI71444.1 |
| 187 | ksb_Benastatin-Hex_12 | 12 | *Streptomyces sp. A2991200* | CAM58799.1 |
| 188 | ksb_Bipentaromycin_10 | 10 | *Streptomyces sp. NRRL F-6131* | WP_030301370.1 |
| 189 | ksb_Brasiliquinone-Pr_10 | 10 | *Nocardia brasiliensis* | WP_042264335.1 |
| 190 | ksb_Calixanthomycin_12 | 12 | *uncultured bacterium* | AJD20009.1 |
| 191 | ksb_Cervimycin-malonoamyl_9 | 9 | *Streptomyces sp. CS113* | OWA01597.1 |
| 192 | ksb_Chartreusin'_10 | 10 | *Streptomyces chartreusis* | AXS67803.1 |
| 193 | ksb_Chartreusin_10 | 10 | *Streptomyces chartreusis* | CAH10162.1 |
| 194 | ksb_Chattamycin_10 | 10 | *Streptomyces lydicus* | WP_046929080.1 |
| 195 | ksb_Chelocardin_10 | 10 | *Amycolatopsis sulphurea* | AHD25927.1 |
| 196 | ksb_Chlortetracycline_10 | 10 | *Kitasatospora aureofaciens* | AEI98665.1 |
| 197 | ksb_Chromomycin_10 | 10 | *Streptomyces griseus subsp. griseus* | CAE17526.1 |
| 198 | ksb_Chrysomycin-Pr_10 | 10 | *Streptomyces albaduncus* | CBH32089.1 |
| 199 | ksb_Cinerubin-Pr_10 | 10 | *Streptomyces sp. SPB074* | EDY42531.1 |
| 200 | ksb_Clostrubin'_X | 10 | *Clostridium puniceum* | WP_077846909.1 |
| 201 | ksb_Clostrubin_X | 10 | *Clostridium beijerinckii* | WP_077837155.1 |
| 202 | ksb_Collinone_13 | 13 | *Streptomyces collinus* | AAG26880.1 |
| 203 | ksb_CosmomycinC-Pr_10 | 10 | *Streptomyces sp. CNT302* | WP_018891747.1 |
| 204 | ksb_Cosmomycin-Pr_10 | 10 | *Streptomyces olindensis* | ABC00728.1 |
| 205 | ksb_Cur_12 | 12 | *Streptomyces cyaneus* | CAA44381.1 |

| | | | | |
|---|---|---|---|---|
| 206 | ksb_Cysgriseusin_10 | 10 | *Streptomyces cyaneofuscatus* | WP_030562169.1 |
| 207 | ksb_Cytorhodin-Pr_10 | 10 | *Streptomyces sp. SCSIO 1666* | ATJ00754.1 |
| 208 | ksb_Dactylocycline_10 | 10 | *Dactylosporangium sp. SC14051* | AFU65895.1 |
| 209 | ksb_Daunorubicin-Pr_10 | 10 | *Streptomyces sp.* | AAA87619.1 |
| 210 | ksb_Dendrubin_9 | 9 | *Dendrosporobacter quercicolus* | WP_092069525.1 |
| 211 | ksb_Doxorubicin'-Pr_10 | 10 | *Streptomyces peucetius* | WP_100109101.1 |
| 212 | ksb_Doxorubicin-Pr_10 | 10 | *Streptomyces peucetius* | AAA65207.1 |
| 213 | ksb_Dutomycin_10 | 10 | *Streptomyces minoensis* | AKD43513.1 |
| 214 | ksb_Elloramycin_10 | 10 | *Streptomyces olivaceus* | CAP12601.1 |
| 215 | ksb_Enduracyclinone_13 | 13 | *Nonomuraea sp.* | AYP71360.1 |
| 216 | ksb_Enterocin-Bn_8 | 8 | *Streptomyces maritimus* | AAF81729.1 |
| 217 | ksb_Erdacin_8 | 8 | *uncultured soil bacterium V167* | ACX83618.1 |
| 218 | ksb_Fasamycin_13 | 13 | *N/A* | AEM44267.1 |
| 219 | ksb_FD-594-Bu_13 | 13 | *Streptomyces sp. TA-0256* | BAJ52682.1 |
| 220 | ksb_Fluostatin_10 | 10 | *uncultured bacterium BAC AB649/1850* | AEE65467.1 |
| 221 | ksb_FluostatinM_10 | 10 | *Micromonospora rosaria* | ALJ99854.1 |
| 222 | ksb_FluostatinMQ_10 | 10 | *Streptomyces albus* | WP_040253445.1 |
| 223 | ksb_Fogacin_8 | 8 | *Actinoplanes missouriensis 431* | BAL90285.1 |
| 224 | ksb_FogacinC-HCS_8 | 8 | *Actinoplanes missouriensis 431* | BAL90262.1 |
| 225 | ksb_Formicamycin_13 | 13 | *Streptomyces sp. KY5* | AQP25564.1 |
| 226 | ksb_Frankiamicin_12 | 12 | *Frankia sp. EAN1pec* | ABW11826.1 |
| 227 | ksb_FredericamycinC_13 | 13 | *Streptomyces albus subsp. chlorinus* | NSC21570.1 |
| 228 | ksb_Fredericamycin-Hex_13 | 13 | *Streptomyces griseus* | AAQ08917.1 |
| 229 | ksb_Frenolicin-Bu_8 | 8 | *Streptomyces roseofulvus* | AAC18108.1 |

| | | | | |
|---|---|---|---|---|
| 230 | ksb_Frigocyclinone_10 | 10 | *Streptomyces griseus* | QDG00825.1 |
| 231 | ksb_Gaudimycin_10 | 10 | *Streptomyces sp. PGA64* | AAK57526.1 |
| 232 | ksb_Gilvocarcin-Pr_10 | 10 | *Streptomyces griseoflavus* | AAP69574.1 |
| 233 | ksb_Granaticin''_8 | 8 | *Streptomyces exfoliatus* | WP_137988252.1 |
| 234 | ksb_Granaticin'_8 | 8 | *Streptomyces violaceoruber* | CAA09654.1 |
| 235 | ksb_Granaticin_8 | 8 | *Streptomyces vietnamensis* | ADO32787.1 |
| 236 | ksb_Grincamycin_10 | 10 | *Streptomyces lusitanus* | AGO50611.1 |
| 237 | ksb_Griseorhodin_13 | 13 | *Streptomyces sp. JP95* | AAM33654.1 |
| 238 | ksb_Griseusin_10 | 10 | *Streptomyces griseus* | CAA54859.1 |
| 239 | ksb_Hatomarubigin_10 | 10 | *Streptomyces sp. 2238-SVT4* | BAJ07844.1 |
| 240 | ksb_Hedamycin-Hex_10 | 10 | *Streptomyces griseoruber* | AAP85361.1 |
| 241 | ksb_Heliquinomycin_13 | 13 | *Streptomyces piniterrae* | WP_136741437.1 |
| 242 | ksb_Hexaricin_13 | 13 | *Streptosporangium sp. FXJ7.131* | AMK51280.1 |
| 243 | ksb_Hiroshidine_8 | 8 | *Streptomyces hiroshimensis* | QBK46636.1 |
| 244 | ksb_Huanglongmycin_9 | 9 | *Streptomyces sp. CB09001* | AXL88815.1 |
| 245 | ksb_Hyaluromycin_13 | 13 | *Streptomyces hyaluromycini* | WP_089099013.1 |
| 246 | ksb_Hydroxyfujianmycin_10 | 10 | *Streptomyces alni* | WP_093715798.1 |
| 247 | ksb_IFQs_8 | 8 | *Streptomyces sp. CB01883* | APR73638.1 |
| 248 | ksb_Isatropolone-Bu_8 | 8 | *Streptomyces sp.* | ARG41912.1 |
| 249 | ksb_Isoindolinomycin-Gly_8 | 8 | *Streptomyces sp. SoC090715LN-16* | BBE36465.1 |
| 250 | ksb_Jadomycin_10 | 10 | *Streptomyces venezuelae ATCC 10712* | AAB36563.1 |
| 251 | ksb_JBIR-76_8 | 8 | *Streptomyces sp. RI-77* | BAU98042.1 |
| 252 | ksb_Julichrome'_9 | 9 | *Streptomyces sampsonii* | QNL10614.1 |
| 253 | ksb_Julichrome_9 | 9 | *Streptomyces afghaniensis* | WP_037671486.1 |

| | | | | |
|---|---|---|---|---|
| 254 | ksb_Keyicin_10 | 10 | *Micromonospora sp. WMMB235* | OHX01517.1 |
| 255 | ksb_Kiamycin_9 | 9 | *Streptomyces sp. W007* | EHM27506.1 |
| 256 | ksb_Kinamycin_10 | 10 | *Streptomyces murayamaensis* | AAO65347.1 |
| 257 | ksb_kinanthraquinone_9 | 9 | *Streptomyces sp. SN-593* | WP_202233565.1 |
| 258 | ksb_Komodoquinone'_10 | 10 | *Streptomyces erythrochromogenes* | WP_031147017.1 |
| 259 | ksb_Komodoquinone_10 | 10 | *Streptomyces sp. NRRL S-378* | WP_030957356.1 |
| 260 | ksb_Kosinostatin_10 | 10 | *Micromonospora sp. TP-A0468* | AFJ52675.1 |
| 261 | ksb_Lactonamycin-Gly_10 | 10 | *Streptomyces rishiriensis* | ABX71115.1 |
| 262 | ksb_LactonamycinZ-Gly_10 | 10 | *Streptomyces sanglieri* | ABX71149.1 |
| 263 | ksb_Landomycin_10 | 10 | *Streptomyces cyanogenus* | AAD13537.1 |
| 264 | ksb_LandomycinE'_10 | 10 | *uncultured bacterium* | AEM44219.1 |
| 265 | ksb_LandomycinE_10 | 10 | *Streptomyces globisporus* | AID46991.1 |
| 266 | ksb_LL-D49194_1-Ile_10 | 10 | *Streptomyces vinaceusdrappus* | QDQ37911.1 |
| 267 | ksb_Lomaiviticin''-Pr_10 | 10 | *Salinispora pacifica* | AHZ61893.1 |
| 268 | ksb_Lomaiviticin'-Pr_10 | 10 | *Salinispora pacifica* | AHF72795.1 |
| 269 | ksb_Lomaiviticin-Pr_10 | 10 | *Salinispora tropica CNB-440* | ABP54675.1 |
| 270 | ksb_Lugdunomycin_10 | 10 | *Streptomyces sp. QL37* | PPQ57490.1 |
| 271 | ksb_Lysolipin_12 | 12 | *Streptomyces tendae* | CAM34344.1 |
| 272 | ksb_Maduralactomycin_10 | 10 | *Actinomadura rubteroloni* | WP_103562971.1 |
| 273 | ksb_Mayamycin_10 | 10 | *Streptomyces sp.* | AVO00814.1 |
| 274 | ksb_Medermycin_8 | 8 | *Streptomyces sp. AM-7161* | BAC79045.1 |
| 275 | ksb_Mensacaricin_10 | 10 | *Streptomyces bottropensis* | AHL46685.1 |
| 276 | ksb_Metamycin-iPr_8_9 | 8/9 | *Rothia dentocariosa* | VEJ30650.1 |
| 277 | ksb_Metathramycin_10 | 10 | *uncultured bacterium* | QVQ68798.1 |

| | | | | |
|---|---|---|---|---|
| 278 | ksb_Methyltetrangomycin_10 | 10 | *Streptomyces sp. Go-475* | WP_114256227.1 |
| 279 | ksb_Mithramycin_10 | 10 | *Streptomyces argillaceus* | CAA61990.1 |
| 280 | ksb_Murayaquinone_10 | 10 | *Streptomyces griseoruber* | AQW35059.1 |
| 281 | ksb_Naphthocyclinone_8 | 8 | *Streptomyces arenae* | AAD20268.1 |
| 282 | ksb_Nenestatin-Pr_10 | 10 | *Micromonospora echinospora* | ARD70902.1 |
| 283 | ksb_Nivetetracyclate-Pr_10 | 10 | *Streptomyces sp. Ls2151* | AGZ78375.1 |
| 284 | ksb_Nocardiopsistin-iPr_10 | 10 | *Nocardiopsis sp.* | QOP59271.1 |
| 285 | ksb_Nogalamycin_10 | 10 | *Streptomyces nogalater* | CAA12018.1 |
| 286 | ksb_Oviedomycin'_10 | 10 | *Streptomyces antibioticus* | ARK36156.1 |
| 287 | ksb_Oviedomycin_10 | 10 | *Streptomyces antibioticus* | CAG14966.1 |
| 288 | ksb_Oxytetracycline'_10 | 10 | *Streptomyces rimosus subsp. rimosus ATCC 10970* | ELQ83293.1 |
| 289 | ksb_Oxytetracycline_10 | 10 | *Streptomyces rimosus* | AAZ78326.1 |
| 290 | ksb_Paramagnetoquinone_X | 10 | *Actinoallomurus sp.* | ASA49565.1 |
| 291 | ksb_PD116740_10 | 10 | *Streptomyces sp. WP 4669* | AAO65363.1 |
| 292 | ksb_Persiamycin_10 | 10 | *Streptomonospora sp. PA3* | WP_156000393.1 |
| 293 | ksb_Piloquinone-Leu_9 | 9 | *Streptomyces sp. FXJ8.102* | QFS19045.1 |
| 294 | ksb_Polyketomycin_10 | 10 | *Streptomyces diastatochromogenes* | ACN64835.1 |
| 295 | ksb_Pradimicin_12 | 12 | *Actinomadura hibisca* | ABM21748.1 |
| 296 | ksb_Pyxidicycline_9 | 9 | *Pyxidicoccus fallax* | AXM42928.1 |
| 297 | ksb_Qinimycin_8 | 8 | *Streptomyces roseifaciens* | WP_058047373.1 |
| 298 | ksb_R1128-Leu_8 | 8 | *Streptomyces sp. R1128* | AAG30188.1 |
| 299 | ksb_Ravidomycin-Pr_10 | 10 | *Streptomyces ravidus* | CBH32808.1 |
| 300 | ksb_Resistomycin_X | 10 | *Streptomyces resistomycificus* | CAE51175.1 |

| | | | | |
|---|---|---|---|---|
| 301 | ksb_Rhodomycin-Pr_10 | 10 | *Streptomyces purpurascens* | WP_189725835.1 |
| 302 | ksb_Rishirilide'-Leu_9 | 9 | *Streptomyces olivaceus* | AWM72903.1 |
| 303 | ksb_Rishirilide-Leu_9 | 9 | *Streptomyces bottropensis* | AHL46709.1 |
| 304 | ksb_Rubrolone-Bu_8 | 8 | *Streptomyces sp. KIB-H033* | AOZ61213.1 |
| 305 | ksb_Rubromycin'_13 | 13 | *Streptomyces collinus* | WP_073806071.1 |
| 306 | ksb_Rubromycin_13 | 13 | *unclassified Streptomyces* | AAG03068.1 |
| 307 | ksb_Saprolmycin_10 | 10 | *Streptomyces sp. TK08046* | BAV17000.1 |
| 308 | ksb_SaquayamycinA_10 | 10 | *Streptomyces sp.* | ARO44656.1 |
| 309 | ksb_SaquayamycinZ_10 | 10 | *Micromonospora sp. Tu 6368* | ACP19354.1 |
| 310 | ksb_Sch_12 | 12 | *Streptomyces halstedii* | AAA02834.1 |
| 311 | ksb_Sch47554_10 | 10 | *Streptomyces sp. SCC 2136* | CAH10116.1 |
| 312 | ksb_Setomimycin_9 | 9 | *Streptomyces aurantiacus* | WP_037658550.1 |
| 313 | ksb_SF2575_10 | 10 | *Streptomyces sp. SF2575* | ADE34519.1 |
| 314 | ksb_Simocyclinone'_10 | 10 | *Streptomyces sp. NRRL B-24484* | WP_030269265.1 |
| 315 | ksb_Simocyclinone_10 | 10 | *Streptomyces antibioticus* | AAK06785.1 |
| 316 | ksb_Steffimycin_10 | 10 | *Streptomyces steffisburgensis* | CAJ42319.1 |
| 317 | ksb_Streptoketide-Pr_10 | 10 | *Streptomyces sp. Tue6314* | QED90612.1 |
| 318 | ksb_Tetarimycin_10 | 10 | *uncultured bacterium* | AFY23043.1 |
| 319 | ksb_Tetracenomycin_10 | 10 | *Streptomyces glaucescens* | AAA67516.1 |
| 320 | ksb_Thioangucycline_10 | 10 | *Streptomyces sp. CB00072* | WP_073867015.1 |
| 321 | ksb_TLN-05220-Ile_13 | 13 | *Micromonospora echinospora subsp. challisensis* | ADB23392.1 |
| 322 | ksb_Trioxacarcin-Ile_10 | 10 | *Streptomyces bottropensis* | AKT74261.1 |
| 323 | ksb_Turbinmicin_13 | 13 | *Micromonospora sp. WMMC415* | WP_155333335.1 |
| 324 | ksb_Urdamycin_10 | 10 | *Streptomyces fradiae* | CAA60570.1 |

| | | | | |
|---|---|---|---|---|
| 325 | ksb_UT-X26_F129_10 | 10 | *Streptomyces aureus* | AYU66234.1 |
| 326 | ksb_Warkmycin_10 | 10 | *Streptomyces sp. CS057* | OWA25244.1 |
| 327 | ksb_Wexrubicin_X | 10 | *Eubacteriales* | WP_025578805.1 |
| 328 | ksb_WhiE_12 | 12 | *Streptomyces coelicolor* | CAA39409.1 |
| 329 | ksb_WS79089D_13 | 13 | *Streptosporangium roseum DSM 43021* | WP_2890797.1 |
| 330 | ksb_X26_10 | 10 | *uncultured bacterium* | AEM44308.1 |
| 331 | ksb_Xantholipin_13 | 13 | *Streptomyces flavogriseus* | ADE22314.1 |
| 332 | ksb_ZHMpentangular_13 | 13 | *Streptomyces viridochromogenes* | WP_003993029.1 |

**Supplementary Table S2.** Index of sequences from Cluster A.

| **Cluster A_Product name** | **ClusterA_Index** |
|---|---|
| FABF | 0 |
| ksa_Abx_13 | 2 |
| ksa_Anthrabenzoxocinone_13 | 10 |
| ksa_Calixanthomycin_12 | 24 |
| ksa_Dutomycin_10 | 47 |
| ksa_Huanglongmycin_9unknown | 78 |
| ksa_LandomycinE'_10unknown | 98 |
| ksa_LandomycinE_10unknown | 99 |
| ksa_Murayaquinone_10unknown | 114 |
| ksa_Oxytetracycline'_10unknown | 122 |
| ksa_R1128-Leu_8unknown | 132 |
| ksa_Rubrolone-Bu_8unknown | 138 |
| ksa_SaquayamycinA_10 | 142 |
| ksa_Sch47554_10unknown | 145 |
| ksa_Simocyclinone'_10unknown | 148 |
| ksb_Piloquinone-Leu_9 | 293 |
| ksb_Pradimicin_12 | 295 |
| ksb_Rubrolone-Bu_8 | 304 |

**Supplementary Table S3.** Clustering Information.

| Cluster | Number of sequences | PKS Sequence index |
|---|---|---|
| KS_3 | 8 | 0, 24, 47, 78, 114, 132, 142, 145 |
| KS_2 | 146 | 1, 3, 4, 5, 6, 7, 8, 9, 12-23, 25-33, 36-43, 45-46, 48-97, 100-109, 111-113, 115-121, 123-131, 133-137, 139-141, 143-144, 146-147, 149-160, 162-166 |
| KS_0 | 7 | 2, 10, 98, 99, 122, 138, 148 |
| KS_4 | 6 | 11, 34, 35, 44, 110, 161 |
| CLF_5 | 53 | 167-169, 174-176, 178, 186, 195, 199, 202, 205, 211-212, 214, 219-220, 226, 229, 233-235, 237-238, 241, 243, 245, 248-249, 271-272, 277, 279, 281, 283, 288-290, 297-298, 302-303, 305-306, 310, 313, 317, 319, 325, 328, 330-332 |
| CLF_1 | 99 | 170-174, 179-198, 203-204, 206-208, 213, 215-218, 221-225, 228, 230-232, 236, 239-240, 242, 244, 246-247, 250-270, 273-274, 278, 280, 282, 284-287, 291-292, 294, 299-301, 307-309, 311-312, 314-315, 318, 320-324, 326, 329 |
| CLF_4 | 6 | 177, 200, 201, 210, 276, 327 |
| CLF_3 | 4 | 209, 227, 275, 316 |
| CLF_0 | 3 | 293, 295, 304 |
| CLF_2 | 1 | 296 |

**Supplementary Table S4.** Edge weights between clusters.

| Edges | Weights |
|---|---|
| CLF0 to KS0 | 1638452.91 |
| KS0 to KS3 | 105408.77 |
| CLF0 to KS2 | 5634.28 |
| CLF0 to CLF5 | 350.86 |
| KS3 to KS3 | 131.33 |
| CLF0 to CLF3 | 128.58 |
| CLF4 to KS4 | 109.02 |
| KS3 to CLF4 | 58.28 |
| KS4 to KS5 | 44.23 |
| CLF5 to CLF1 | 41.93 |
| CLF4 to KS2 | 26.98 |
| KS4 to KS2 | 21.07 |
| KS3 to KS2 | 18.38 |
| CLF0 to KS3 | 17.22 |
| CLF2 to KS2 | 14.00 |
| CLF4 to KS1 | 10.36 |
| KS3 to KS4 | 8.38 |
| CLF4 to CLF2 | 4.27 |
| KS0 to CLF3 | 2.31 |
| KS3 to CLF1 | 2.00 |
| CLF4 to CLF5 | 1.00 |
| CLF1 to KS2 | 1.00 |

**Supplementary Table S5.** Index of Non-oxidative PKS sequences.

| Non-oxidative PKS product | KS_index | CLF_index |
|---|---|---|
| AQ-256_8 | 11 | 177 |
| Clostrubin'_X | 34 | 200 |
| Clostrubin_X | 35 | 201 |
| Dendrubin_9 | 44 | 210 |
| Pyxidicycline_9 | 130 | 296 |
| Resistomycin_X | 134 | 300 |
| Wexrubicin_X | 161 | 327 |

**Supplementary Table S6.** DeepFRI function predictions. Due to space limitations, it is presented in a separate table.

**Supplementary Table S7.** Vectorized DeepFRI protein function prediction. Due to space limitations, it is presented in a separate table.

**Supplementary Table S8.** Pairwise protein function similarity comparison matrix. Due to space limitations, it is presented in a separate table.

**Supplementary Table S9.** Co-evolution distances to ancestral node, the ancestral node position was determined by averaging all vectors in Cluster A.

| **KS / CLF pair** | **KS to Cluster A distance** | **CLF to Cluster A distance** | **KS / CLF distance ratio** |
|---|---|---|---|
| KS 1 / CLF 167 | 0.18 | 0.17 | 1.09 |
| KS 3 / CLF 169 | 0.18 | 0.18 | 1.01 |
| KS 4 / CLF 170 | 0.17 | 0.19 | 0.89 |
| KS 5 / CLF 171 | 0.18 | 0.18 | 0.99 |
| KS 6 / CLF 172 | 0.18 | 0.18 | 1.00 |
| KS 7 / CLF 173 | 0.19 | 0.18 | 1.03 |
| KS 8 / CLF 174 | 0.18 | 0.18 | 1.04 |
| KS 9 / CLF 175 | 0.18 | 0.18 | 1.00 |
| KS 11 / CLF 177 | 0.21 | 0.22 | 0.95 |
| KS 12 / CLF 178 | 0.18 | 0.18 | 0.98 |
| KS 13 / CLF 179 | 0.18 | 0.18 | 1.02 |
| KS 14 / CLF 180 | 0.18 | 0.18 | 1.02 |
| KS 15 / CLF 181 | 0.18 | 0.18 | 0.99 |
| KS 16 / CLF 182 | 0.18 | 0.18 | 1.00 |
| KS 17 / CLF 183 | 0.18 | 0.19 | 0.96 |
| KS 18 / CLF 184 | 0.18 | 0.18 | 1.01 |
| KS 19 / CLF 185 | 0.18 | 0.18 | 1.01 |
| KS 20 / CLF 186 | 0.18 | 0.18 | 1.00 |
| KS 21 / CLF 187 | 0.18 | 0.16 | 1.12 |
| KS 22 / CLF 188 | 0.18 | 0.18 | 1.02 |
| KS 23 / CLF 189 | 0.18 | 0.19 | 0.96 |
| KS 25 / CLF 191 | 0.18 | 0.19 | 0.96 |
| KS 26 / CLF 192 | 0.18 | 0.18 | 0.96 |
| KS 27 / CLF 193 | 0.17 | 0.18 | 0.95 |
| KS 28 / CLF 194 | 0.18 | 0.18 | 0.96 |

| | | | |
|---|---|---|---|
| KS 29 / CLF 195 | 0.18 | 0.18 | 1.02 |
| KS 30 / CLF 196 | 0.18 | 0.19 | 0.95 |
| KS 31 / CLF 197 | 0.18 | 0.18 | 1.00 |
| KS 32 / CLF 198 | 0.18 | 0.18 | 1.00 |
| KS 33 / CLF 199 | 0.18 | 0.18 | 0.99 |
| KS 34 / CLF 200 | 0.22 | 0.22 | 1.01 |
| KS 35 / CLF 201 | 0.22 | 0.21 | 1.02 |
| KS 36 / CLF 202 | 0.18 | 0.18 | 0.98 |
| KS 37 / CLF 203 | 0.18 | 0.18 | 0.98 |
| KS 38 / CLF 204 | 0.18 | 0.18 | 1.00 |
| KS 39 / CLF 205 | 0.18 | 0.18 | 0.97 |
| KS 40 / CLF 206 | 0.18 | 0.18 | 1.00 |
| KS 41 / CLF 207 | 0.18 | 0.18 | 0.98 |
| KS 42 / CLF 208 | 0.18 | 0.18 | 0.96 |
| KS 43 / CLF 209 | 0.18 | 0.13 | 1.38 |
| KS 44 / CLF 210 | 0.21 | 0.2 | 1.02 |
| KS 45 / CLF 211 | 0.18 | 0.18 | 1.03 |
| KS 46 / CLF 212 | 0.18 | 0.18 | 1.03 |
| KS 48 / CLF 214 | 0.18 | 0.18 | 1.02 |
| KS 49 / CLF 215 | 0.18 | 0.18 | 1.01 |
| KS 50 / CLF 216 | 0.18 | 0.18 | 0.99 |
| KS 51 / CLF 217 | 0.18 | 0.17 | 1.05 |
| KS 52 / CLF 218 | 0.18 | 0.18 | 1.02 |
| KS 53 / CLF 219 | 0.18 | 0.18 | 1.00 |
| KS 54 / CLF 220 | 0.18 | 0.19 | 0.97 |
| KS 55 / CLF 221 | 0.18 | 0.19 | 0.95 |
| KS 56 / CLF 222 | 0.18 | 0.18 | 0.96 |
| KS 57 / CLF 223 | 0.18 | 0.17 | 1.02 |
| KS 58 / CLF 224 | 0.18 | 0.17 | 1.03 |
| KS 59 / CLF 225 | 0.18 | 0.18 | 0.98 |
| KS 60 / CLF 226 | 0.18 | 0.18 | 1.03 |
| KS 61 / CLF 227 | 0.18 | 0.15 | 1.24 |
| KS 62 / CLF 228 | 0.18 | 0.18 | 0.99 |
| KS 63 / CLF 229 | 0.18 | 0.18 | 0.97 |
| KS 64 / CLF 230 | 0.18 | 0.18 | 0.98 |
| KS 65 / CLF 231 | 0.18 | 0.19 | 0.98 |
| KS 66 / CLF 232 | 0.18 | 0.18 | 0.99 |
| KS 67 / CLF 233 | 0.18 | 0.18 | 0.99 |
| KS 68 / CLF 234 | 0.18 | 0.17 | 1.04 |
| KS 69 / CLF 235 | 0.18 | 0.18 | 1.03 |
| KS 70 / CLF 236 | 0.18 | 0.18 | 1.00 |
| KS 71 / CLF 237 | 0.18 | 0.18 | 1.01 |
| KS 72 / CLF 238 | 0.18 | 0.18 | 1.00 |

| | | | |
|---|---|---|---|
| KS 73 / CLF 239 | 0.18 | 0.18 | 0.96 |
| KS 74 / CLF 240 | 0.18 | 0.18 | 0.95 |
| KS 75 / CLF 241 | 0.18 | 0.18 | 1.01 |
| KS 76 / CLF 242 | 0.18 | 0.18 | 1.01 |
| KS 77 / CLF 243 | 0.19 | 0.18 | 1.04 |
| KS 79 / CLF 245 | 0.18 | 0.18 | 1.00 |
| KS 80 / CLF 246 | 0.18 | 0.19 | 0.98 |
| KS 81 / CLF 247 | 0.18 | 0.18 | 1.02 |
| KS 82 / CLF 248 | 0.18 | 0.18 | 0.98 |
| KS 83 / CLF 249 | 0.18 | 0.18 | 0.96 |
| KS 84 / CLF 250 | 0.18 | 0.18 | 0.98 |
| KS 85 / CLF 251 | 0.18 | 0.18 | 1.03 |
| KS 86 / CLF 252 | 0.18 | 0.18 | 1.00 |
| KS 87 / CLF 253 | 0.18 | 0.18 | 1.00 |
| KS 88 / CLF 254 | 0.18 | 0.18 | 1.01 |
| KS 89 / CLF 255 | 0.18 | 0.18 | 0.97 |
| KS 90 / CLF 256 | 0.18 | 0.19 | 0.98 |
| KS 91 / CLF 257 | 0.18 | 0.18 | 0.99 |
| KS 92 / CLF 258 | 0.18 | 0.19 | 0.96 |
| KS 93 / CLF 259 | 0.18 | 0.19 | 0.96 |
| KS 94 / CLF 260 | 0.18 | 0.19 | 0.97 |
| KS 95 / CLF 261 | 0.18 | 0.18 | 1.00 |
| KS 96 / CLF 262 | 0.19 | 0.18 | 1.02 |
| KS 97 / CLF 263 | 0.18 | 0.18 | 0.99 |
| KS 100 / CLF 266 | 0.17 | 0.18 | 0.98 |
| KS 101 / CLF 267 | 0.18 | 0.19 | 0.96 |
| KS 102 / CLF 268 | 0.18 | 0.19 | 0.98 |
| KS 103 / CLF 269 | 0.18 | 0.19 | 0.97 |
| KS 104 / CLF 270 | 0.18 | 0.18 | 0.97 |
| KS 105 / CLF 271 | 0.18 | 0.18 | 0.97 |
| KS 106 / CLF 272 | 0.18 | 0.18 | 1.00 |
| KS 107 / CLF 273 | 0.18 | 0.19 | 0.97 |
| KS 108 / CLF 274 | 0.18 | 0.18 | 0.98 |
| KS 109 / CLF 275 | 0.18 | 0.13 | 1.44 |
| KS 110 / CLF 276 | 0.2 | 0.19 | 1.04 |
| KS 111 / CLF 277 | 0.19 | 0.18 | 1.02 |
| KS 112 / CLF 278 | 0.18 | 0.18 | 0.99 |
| KS 113 / CLF 279 | 0.18 | 0.18 | 0.99 |
| KS 115 / CLF 281 | 0.18 | 0.18 | 1.05 |
| KS 116 / CLF 282 | 0.18 | 0.19 | 0.98 |
| KS 117 / CLF 283 | 0.18 | 0.18 | 0.99 |
| KS 118 / CLF 284 | 0.18 | 0.18 | 0.99 |
| KS 119 / CLF 285 | 0.18 | 0.18 | 1.02 |

| | | | |
|---|---|---|---|
| KS 120 / CLF 286 | 0.18 | 0.19 | 0.98 |
| KS 121 / CLF 287 | 0.19 | 0.19 | 1.00 |
| KS 123 / CLF 289 | 0.18 | 0.18 | 1.01 |
| KS 124 / CLF 290 | 0.18 | 0.17 | 1.04 |
| KS 125 / CLF 291 | 0.18 | 0.18 | 0.98 |
| KS 126 / CLF 292 | 0.18 | 0.18 | 1.03 |
| KS 127 / CLF 293 | 0.18 | 0.23 | 0.79 |
| KS 128 / CLF 294 | 0.18 | 0.18 | 0.99 |
| KS 129 / CLF 295 | 0.18 | 0.11 | 1.65 |
| KS 130 / CLF 296 | 0.19 | 0.18 | 1.04 |
| KS 131 / CLF 297 | 0.18 | 0.18 | 1.03 |
| KS 133 / CLF 299 | 0.18 | 0.18 | 1.01 |
| KS 134 / CLF 300 | 0.19 | 0.18 | 1.06 |
| KS 135 / CLF 301 | 0.18 | 0.18 | 1.01 |
| KS 136 / CLF 302 | 0.18 | 0.19 | 0.96 |
| KS 137 / CLF 303 | 0.18 | 0.18 | 0.97 |
| KS 139 / CLF 305 | 0.18 | 0.18 | 0.97 |
| KS 140 / CLF 306 | 0.18 | 0.18 | 0.98 |
| KS 141 / CLF 307 | 0.18 | 0.18 | 0.99 |
| KS 143 / CLF 309 | 0.18 | 0.18 | 0.99 |
| KS 144 / CLF 310 | 0.18 | 0.18 | 1.01 |
| KS 146 / CLF 312 | 0.18 | 0.19 | 0.98 |
| KS 147 / CLF 313 | 0.18 | 0.18 | 1.00 |
| KS 149 / CLF 315 | 0.18 | 0.19 | 0.97 |
| KS 150 / CLF 316 | 0.18 | 0.13 | 1.38 |
| KS 151 / CLF 317 | 0.18 | 0.18 | 0.95 |
| KS 152 / CLF 318 | 0.18 | 0.18 | 0.99 |
| KS 153 / CLF 319 | 0.18 | 0.18 | 1.02 |
| KS 154 / CLF 320 | 0.18 | 0.18 | 0.98 |
| KS 155 / CLF 321 | 0.18 | 0.18 | 1.01 |
| KS 156 / CLF 322 | 0.18 | 0.18 | 0.96 |
| KS 157 / CLF 323 | 0.18 | 0.18 | 0.98 |
| KS 158 / CLF 324 | 0.18 | 0.18 | 0.99 |
| KS 159 / CLF 325 | 0.18 | 0.18 | 1.03 |
| KS 160 / CLF 326 | 0.18 | 0.18 | 0.98 |
| KS 161 / CLF 327 | 0.22 | 0.21 | 1.01 |
| KS 162 / CLF 328 | 0.18 | 0.18 | 0.97 |
| KS 163 / CLF 329 | 0.18 | 0.18 | 1.01 |
| KS 164 / CLF 330 | 0.19 | 0.18 | 1.04 |
| KS 165 / CLF 331 | 0.18 | 0.18 | 0.99 |
| KS 166 / CLF 332 | 0.18 | 0.18 | 0.97 |

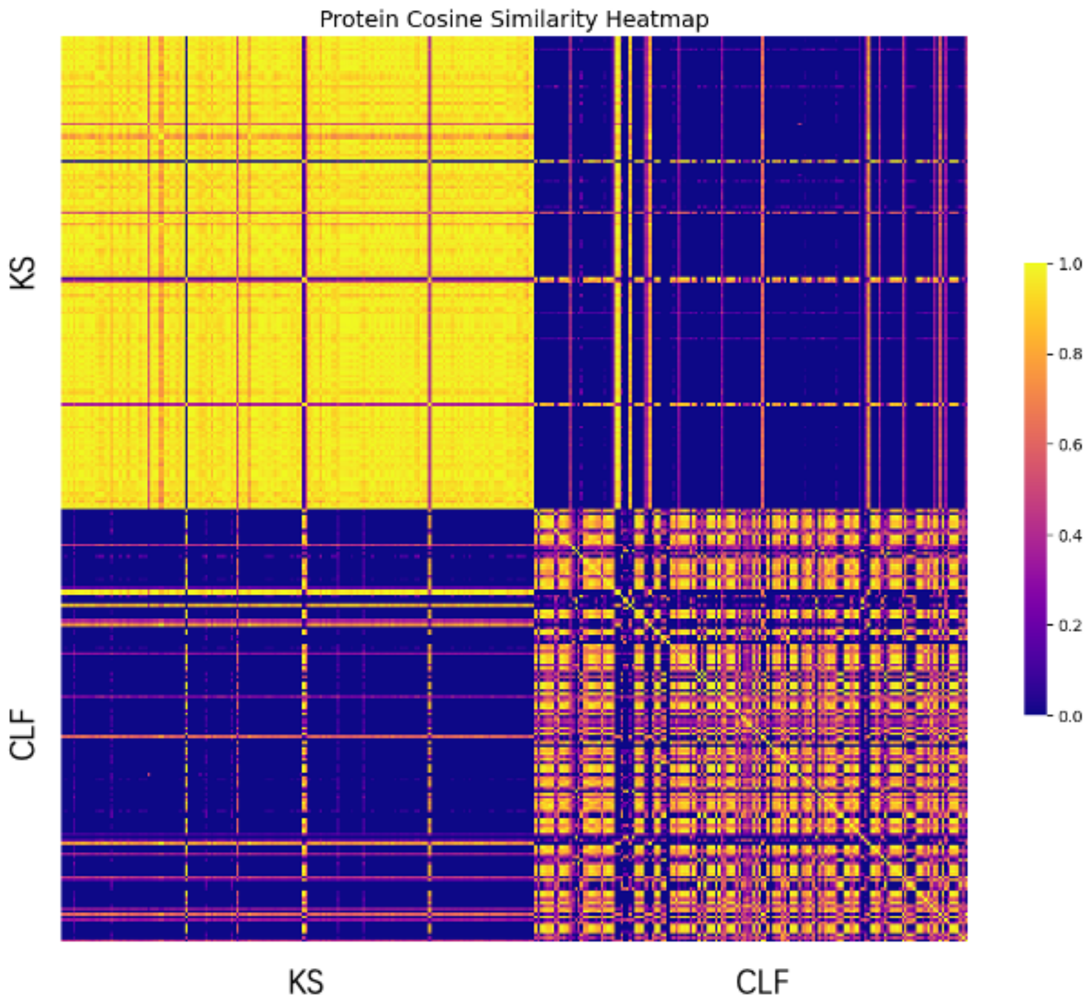


### 2. ESI Figures

**Supplementary Figure S1.** Visualization of DeepFRI predicted function cosine similarities from all 166 pair of T2 PKS.

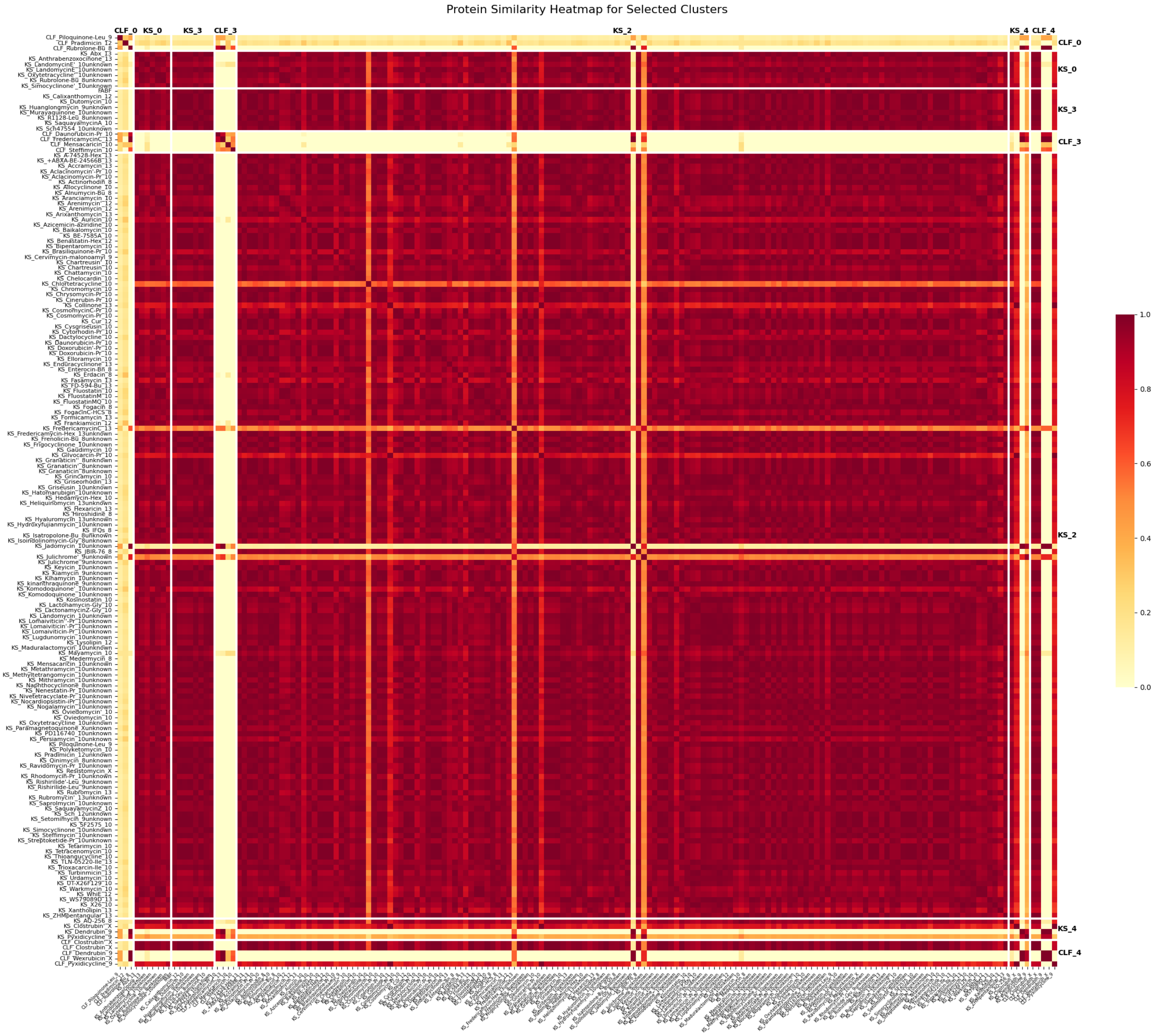


**Supplementary Figure S2.** Visualization of DeepFRI predicted function cosine similarities between selected clusters: CLF_0, KS_0, KS_3, CLF_3, KS_2, KS_4, CLF_4. KS_2 is the majority KS cluster.

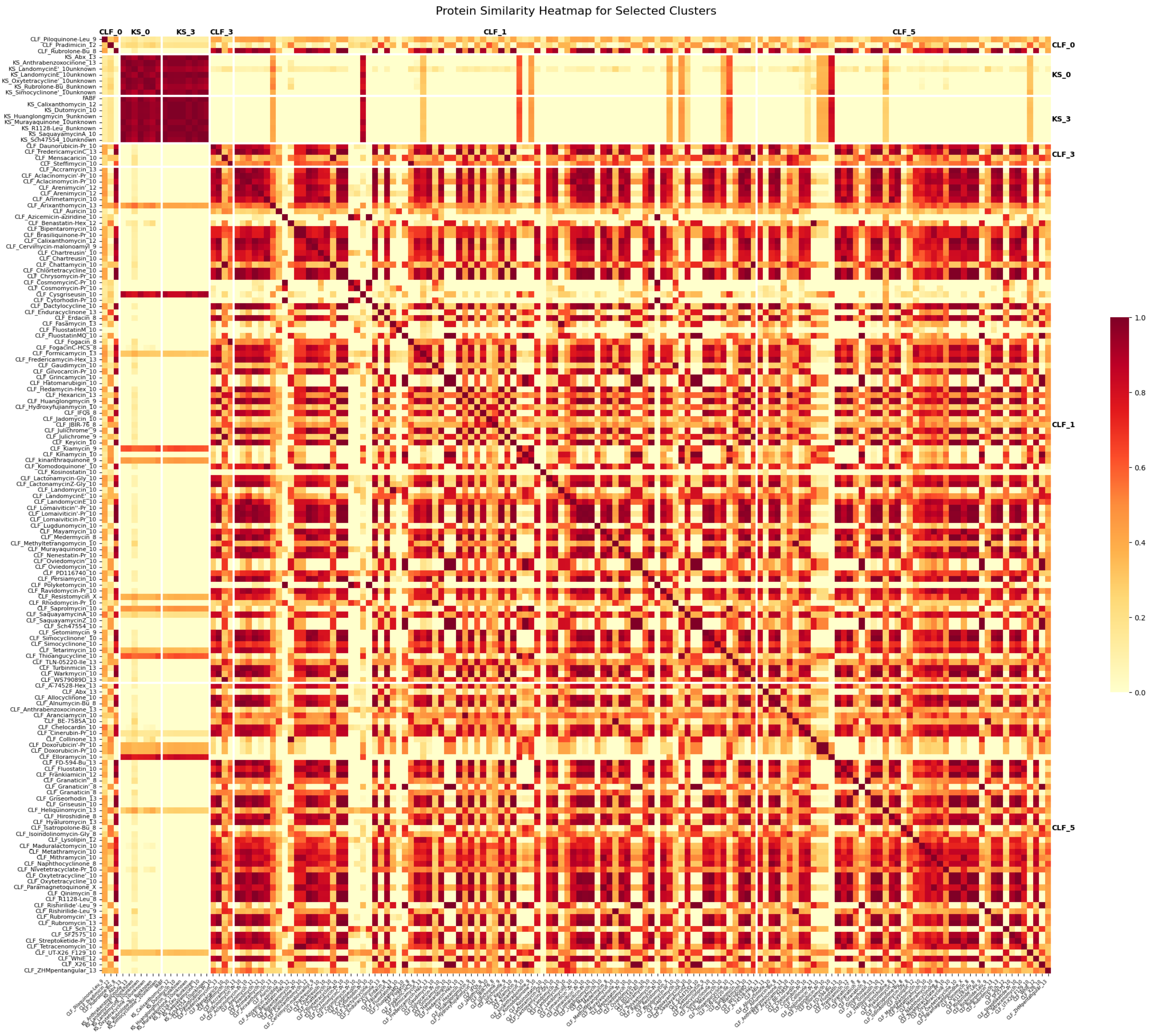


**Supplementary Figure S3.** Visualization of DeepFRI predicted function cosine similarities between selected clusters: CLF_0, KS_0, KS_3, CLF_3, CLF_1, CLF_5. CLF_1 and CLF_5 are the majority CLF clusters.
